# Sex differences in aggression and its neural substrate in a cichlid fish

**DOI:** 10.1101/2023.10.18.562975

**Authors:** Lillian R. Jackson, Mariam Dumitrascu, Beau A. Alward

**Affiliations:** University of Houston, Department of Psychology; University of Houston, Department of Biology and Biochemistry

**Keywords:** aggression, sex differences, sexual dimorphism, neural activation, social behavior network

## Abstract

Aggression is ubiquitous among social species and functions to maintains social dominance hierarchies. The African cichlid fish *Astatotilapia burtoni* is an ideal study species for studying aggression due to their unique and flexible dominance hierarchy. However, female aggression in this species and the neural mechanisms of aggression in both sexes is not well understood. To further understand the potential sex differences in aggression in this species, we characterized aggression in male and female *A. burtoni* in a mirror assay. We then quantified neural activation patterns in brain regions of the social behavior network (SBN) to investigate if differences in behavior are reflected in the brain with immunohistochemistry by detecting the phosphorylated ribosome marker phospho-S6 ribosomal protein (pS6), a marker for neural activation. We found that *A. burtoni* perform both identical and sex-specific aggressive behaviors in response to a mirror assay. We observed sex differences in pS6 immunoreactivity in the Vv, a homolog of the lateral septum in mammals. Males but not females had higher ps6 immunoreactivity in the ATn after the aggression assay. The ATn is a homolog of the ventromedial hypothalamus in mammals, which is strongly implicated in the regulation of aggression in males. Several regions also have higher pS6 immunoreactivity in negative controls than fish exposed to a mirror, implicating a role for inhibitory neurons in suppressing aggression until a relevant stimulus is present. Male and female *A. burtoni* display both similar and sexually dimorphic behavioral patterns in aggression in response to a mirror assay. There are also sex differences in the corresponding neural activation patterns in the SBN. In mirror males but not females, the ATn clusters with the POA, revealing a functional connectivity of these regions that is triggered in an aggressive context in males. These findings suggest that distinct neural circuitry underlie aggressive behavior in male and female *A. burtoni*, serving as a foundation for future work investigating the molecular and neural underpinnings of sexually dimorphic behaviors in this species to reveal fundamental insights into understanding aggression.

## Introduction

Aggression is a complex social behavior defined as agonistic acts directed towards conspecifics and is used to resolve conflict, maintain social dominance hierarchies, gain access to mates, and defend resources like shelter and food ^1,2^. Although aggression is metabolically costly and oftentimes risky, it exists in nearly all species ^3^. Extensive knowledge has been gained on the molecular and neural mechanisms of aggression in both sexes; however, male intrasexual aggression has been studied more extensively in this regard ^3–6^. Moreover, our most in-depth understanding of the mechanisms of aggression come from a few traditional model organisms such as mice, rats, and fruit flies ^3,3,7^. To gain a comprehensive understanding of the molecular and neural control of sexually dimorphic behaviors, models are needed in which 1) both sexes perform aggression in similar contexts and, if possible, 2) both sexes perform the same behaviors in those similar contexts ^8^. Investigating the same behavior in both sexes allows us to identify the fundamental factors influencing aggression and guide research to determine which organizational impacts on the brain shapes the neural and molecular basis of social behaviors adult behavior. However, these goals have been challenging to achieve in current models for studying the molecular and neural control of aggression.

The African cichlid fish *Astatotilapia burtoni* may be an especially useful species in which to gain fundamental insights into the sex differences in aggression and its molecular and neural control ^9–14^. *A. burtoni* exhibit dynamic social interactions and decades of field and laboratory studies have revealed novel insights into the mechanisms governing social behavior. Male *A. burtoni* exist in two dominance states that can be transitioned between depending on the social environment ^15^. Dominant males are brightly colored, have high levels of circulating sex steroid hormones (androgens and estrogens), and perform reproductive and aggressive behaviors. Aggressive behaviors such as attacks and lateral displays are distinct and easily identifiable in behavioral assays (Figure 1). In contrast, subordinate males are drably colored, have low levels of circulating sex steroid hormones, and perform submissive behaviors ^16,17^.

**Figure 1.**
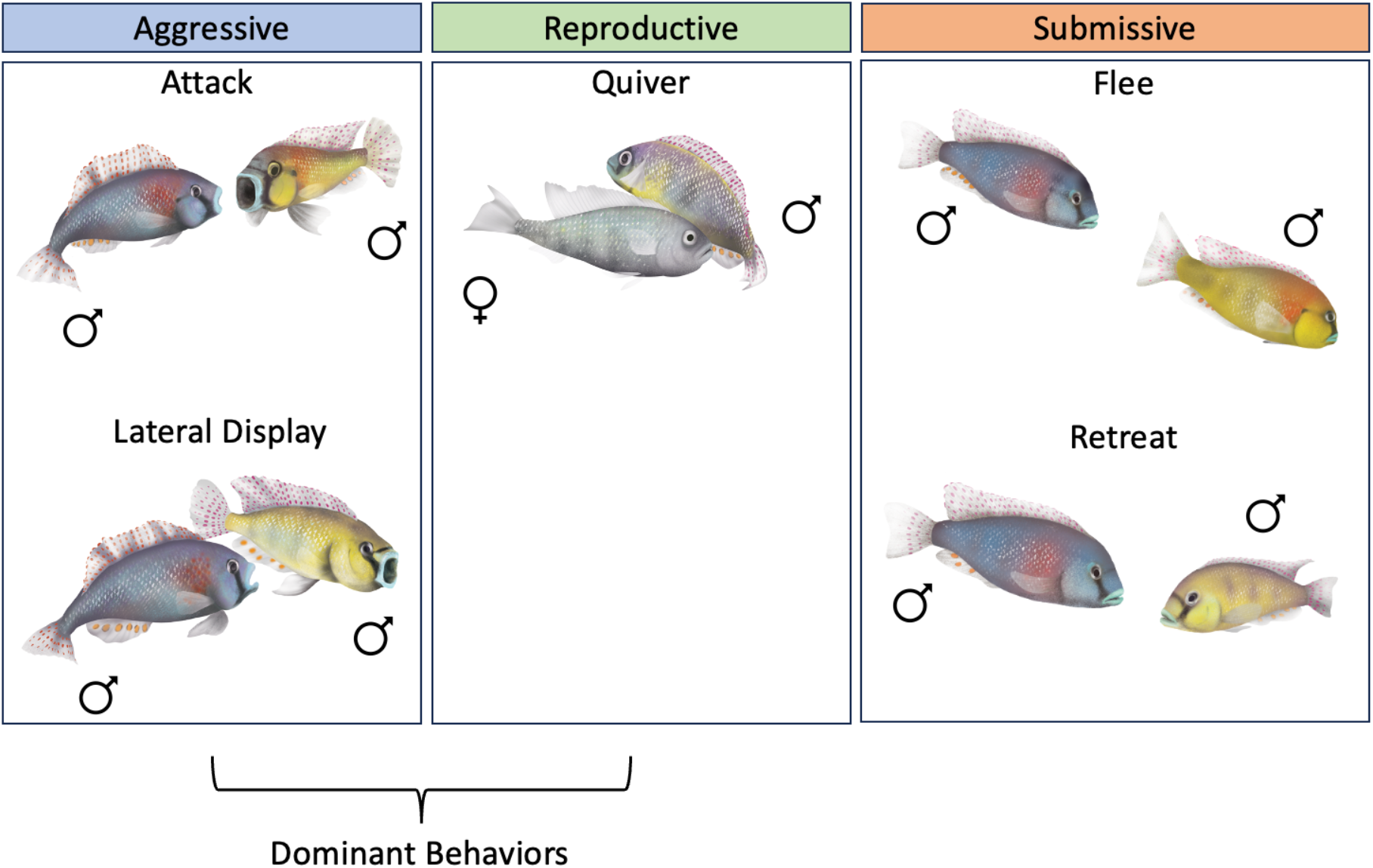
Typical behaviors characterized in *A. burtoni*. Aggressive and reproductive behaviors are typically performed by dominant males. Submissive behaviors are performed by subordinate males.

Female *A. burtoni* do not naturally form a social hierarchy and typically shoal with other females. However, when in all-female communities, females acquire certain male-typical dominance traits including enhanced aggression ^18^, suggesting some mechanisms regulating male-typical dominance behaviors are present in females. Much like the dominance hierarchy in male *A. burtoni*, female *A. burtoni* dominance is plastic and reversible, with dominant females directing aggressive behaviors like chasing and lateral displays towards subordinate females ^18^. Still, outside of all-female communities, aggressive behavioral patterns in *female A. burtoni* have not been characterized. Aggression in both sexes has, however, been studied in two teleost species that have been studied for non-breeding territorial aggression, aggression does not differ between males and females. *Gymnotus omarorum* males and females show no differences in aggressive behaviors ^19,20^ and in *Stegastes nigricans*, the level of aggression does not differ between males and females but territorial behavior differs during the reproductive period ^21,22^, in which females frequently leave their territories but males remain in theirs to guard eggs.

To definitively determine which behaviors *A. burtoni* females use during aggressive interactions, and whether they are similar to male aggressive behaviors, requires assays in which aggression is displayed by both sexes that is readily observable and quantifiable. Here, we characterize aggressive behaviors in male and female *A. burtoni* using a mirror assay. In *A. burtoni* and other model organisms like zebrafish, mirrors elicit aggressive behavioral responses because the fish treats its reflection as another fish ^23,24^. We reasoned that mirror assays would be especially useful for studying aggression in *A. burtoni* since males alter their aggression if an intruder male is larger, smaller, or the same size (within 5% of other males) in length ^25^. Therefore, to successfully compare aggression in males and females requires the use of a paradigm like the mirror assay, which controls for size given that their reflection should be treated as a fish identical in size to it. However, it is important to note the evidence that cichlids and other fish do not treat a mirror exactly like a live opponent including behavioral responses and neural patterns ^24,26^. We argue that since mirrors are able to elicit robust behavioral responses in *A. burtoni* and we are interested in the mechanisms of the behavioral acts themselves, that it is appropriate to use a mirror assay in this study.

In fish included in the mirror assay, we also investigated neural activation patterns as measured by phosphorylated ribosomes throughout brain regions of the social behavior network (SBN) and the social decision-making network (SDMN), two conserved, interconnected circuits critical for integrating social cues and modulating social behaviors ^27,28^. We hypothesized that (1) males and females would perform sexually dimorphic aggressive behaviors during a mirror assay and that these differences would be reflected as (2) sexually dimorphic neural activation patterns. Our findings show suites of convergent and divergent behavioral and neural sex differences in the control of aggression in *A. burtoni*, suggesting independent mechanisms mediate sexually dimorphic behaviors in this species. These results highlight the importance of studying sex differences in diverse organisms to illuminate the fundamental mechanisms underlying the generation of social behavior and suggest *A. burtoni* may be a useful model for understanding the molecular and neural control of sexually dimorphic aggression.

## Methods

### Animal Subjects

Adult *A. burtoni* were bred from a wild-caught stock that originated from Lake Tanganyika, Africa and housed in environmental conditions that mimic their natural equatorial habitat (28 °C; pH 8.0; 12:12 h light/dark cycle with full spectrum illumination; constant aeration). Aquaria contained gravel-covered bottoms with terra cotta pots cut in half to serve as shelters. Fish were fed cichlid flakes daily. All experimental procedures were approved by the University of Houston Institutional Animal Care and Use Committee (Protocol #202000001).

### Mirror Assay behavior recording

Male (n=8) and female (n=9) *Astatotilapia burtoni* were housed in a pre-assay tank for approximately two weeks prior to being assayed in a mirror assay. The pre-assay tank consisted of a 60.5 L plexiglass tank with a clear perforated divider that separated males and females on separate halves to allow males and females to interact but prevent them from mating. Subjects were housed in groups of 6 fish at a time, including 3 gravid females and 3 dominant males on each side. After two weeks in the pre-assay tank, each subject’s behaviors were recorded in a mirror assay. One dominant male and one female were assayed each day until all the subjects in a pre-assay group were assayed (Figure 2).

**Figure 2.**
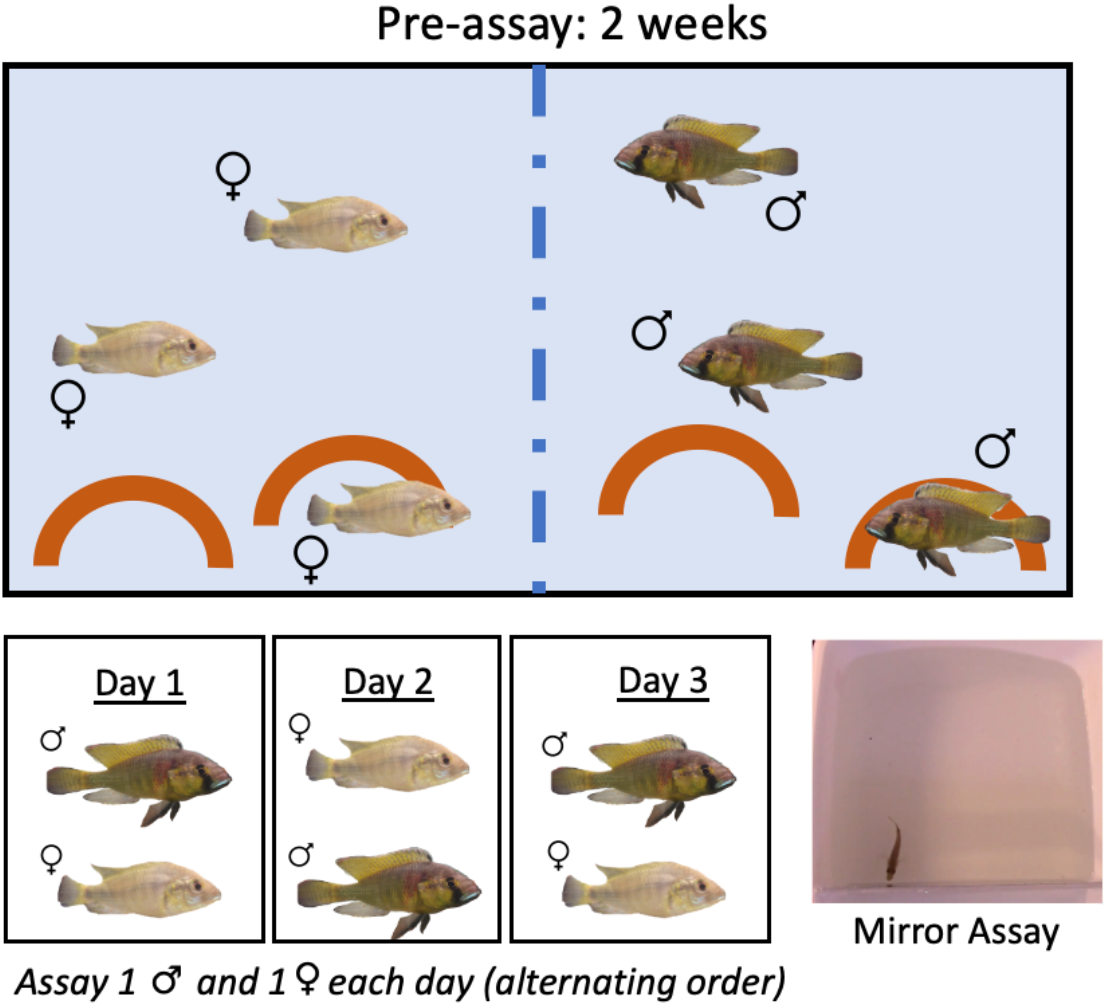
Pre-assay schematic for negative control and mirror fish. Males and females were housed in a tank separated by a perforated divider for 2 weeks. Following this pre-assay period, males and females were assayed in pairs on consecutive days in a mirror assay.

The mirror assay tank consisted of a white tub (Sterilite; 400 mm x 318 mm x 12 mm) filled 6 cm deep with UV-sterilized aquaria water with a mirror replacing one side of the assay tank. An opaque cover was first placed on the mirror for a 15-minute habituation period to allow subjects to acclimate to the assay tank. The opaque cover was then removed, and 30 minutes of behavior was recorded. We also included negative control males (n= 5) and females (n= 8) in the same exact pre-assay conditions except the mirror cover was not lifted after the 15-minute habituation period. Immediately following the mirror assay, subjects were euthanized in an ice bath for 1-2 minutes before rapid cervical transection followed immediately by the harvesting of tissue.

### Behavioral analysis

Behavior was recorded using a digital video camera and was quantified using the freely available BORIS (Behavioral Observation Research Interactive Software). Multiple types of behavior were quantified: behaviors considered not to be aggressive or “neutral” (tap, graze); subordinate behaviors (flee, retreat); aggressive behaviors (attack, lateral display, rostral display); and a male-typical reproductive behavior (quiver) (Figure 1). Taps were defined as making gentle mouth contact with the mirror. Grazing was defined as swimming alongside the mirror for at least 3 seconds. Fleeing was defined as an abrupt turn and swim away from the mirror. Retreats were defined as a rapid retreat swim from the mirror. Attacks were defined as a rapid swim towards the mirror, making mouth contact, and vigorously wiggling the body. Lateral displays were defined as the presentation of the side of the body towards the mirror, conforming the body at times in a slightly convex shape while lateral to the mirror with erect fins. Lateral displays were further separated into type 1 and type 2 lateral displays to distinguish between more and less vigorous lateral displays respectively. Rostral displays were defined as flaring the opercula towards the mirror without making direct contact with the mirror. Quivers were defined as a rapid vibration of the body towards the mirror while conforming in a convex shape. A group of behaviors were summed together for their mouth movements, termed mouth contact (summed attacks, taps, and rostral displays). Additionally, total behaviors were calculated as the sum of all behaviors and total aggression was the sum of all aggressive behaviors (attacks, lateral displays, rostral displays, and quivers). Furthermore, another behavior was identified that appears to be associated with aggression that we have termed an s-curve display, defined as a curved body conformation facing the mirror at a distance.

Tracking and activity data was collected on pre-mirror and mirror assay videos using Noldus EthoVisionXT software. Center-point detection was used to track the subjects throughout the assay tank. Distance moved (cm) was calculated as the accumulated distance the subject moved throughout the assay duration. Velocity (cm/s) was calculated as distance over time. Activity was measured as the average % of pixels changed over time. We defined a zone close to the mirror to measure the proximity of subjects near the mirror, extending 2.4 cm out from the mirror. Zone duration (s) was calculated as the cumulative duration of time spent in the zone. Zone frequency was calculated as the number of times the center-point of the subject entered the zone. Zone latency was defined as the lapse of time until the zone is first entered. Total and mean distances to the zone (cm) were calculated as the cumulative and mean distances from the zone respectively.

**Table 1.**
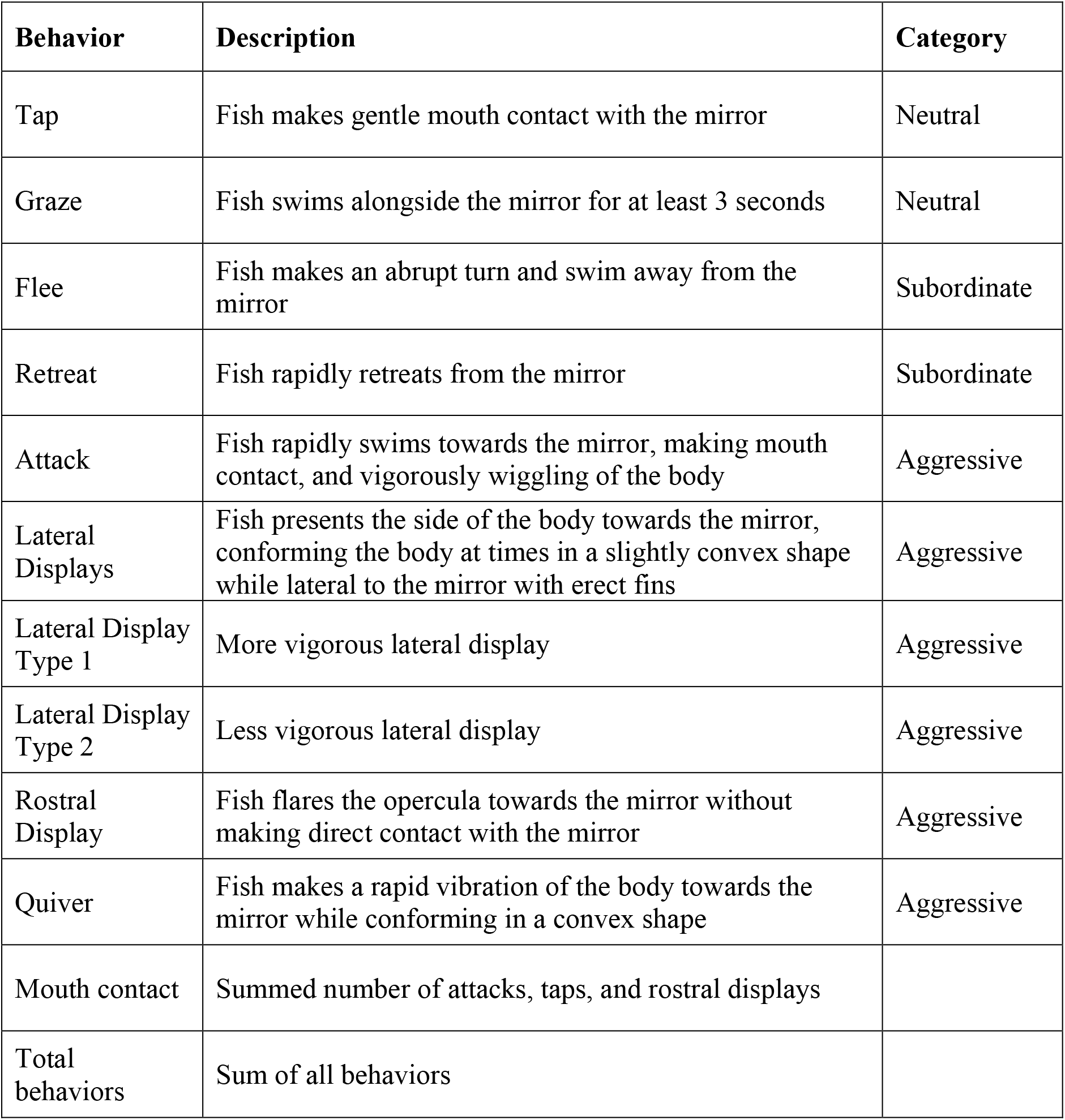
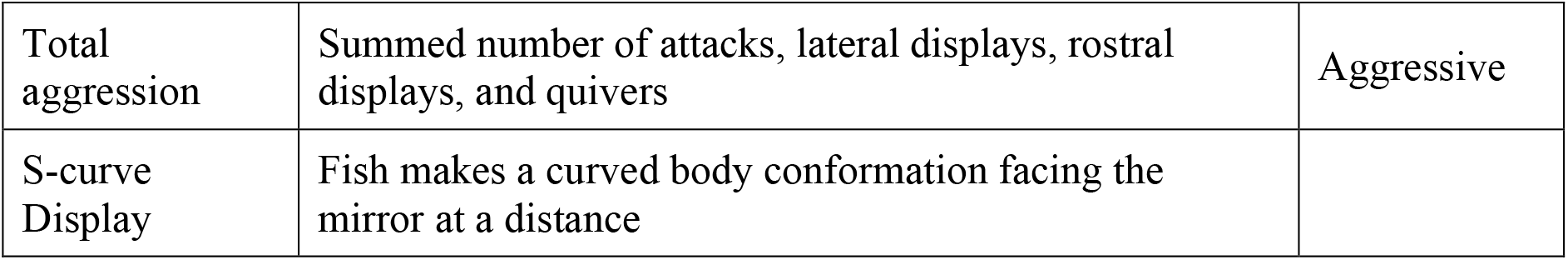
Ethogram used to score behaviors in BORIS.

| **Behavior** | **Description** | **Category** |
| --- | --- | --- |
| Tap | Fish makes gentle mouth contact with the mirror | Neutral |
| Graze | Fish swims alongside the mirror for at least 3 seconds | Neutral |
| Flee | Fish makes an abrupt turn and swim away from the mirror | Subordinate |
| Retreat | Fish rapidly retreats from the mirror | Subordinate |
| Attack | Fish rapidly swims towards the mirror, making mouth contact, and vigorously wiggling of the body | Aggressive |
| Lateral Displays | Fish presents the side of the body towards the mirror, conforming the body at times in a slightly convex shape while lateral to the mirror with erect fins | Aggressive |
| Lateral Display Type 1 | More vigorous lateral display | Aggressive |
| Lateral Display Type 2 | Less vigorous lateral display | Aggressive |
| Rostral Display | Fish flares the opercula towards the mirror without making direct contact with the mirror | Aggressive |
| Quiver | Fish makes a rapid vibration of the body towards the mirror while conforming in a convex shape | Aggressive |
| Mouth contact | Summed number of attacks, taps, and rostral displays |  |
| Total behaviors | Sum of all behaviors |  |
| Total aggression | Summed number of attacks, lateral displays, rostral displays, and quivers | Aggressive |
| S-curve Display | Fish makes a curved body conformation facing the mirror at a distance |  |

A series of dyad assays similar to that used recently in male *A. burtoni* ^25^ were also performed to ensure that female aggressive behaviors observed in the mirror assay were also performed towards a live opponent. Gravid resident females (n = 5) were housed in a 20.8 L glass tank containing gravel, half of a terra cotta pot, and constant aeration for two days before being introduced to a gravid intruder female. The intruder female was an unfamiliar conspecific that was size-matched within 5% of the resident’s standard length. After the intruder female was added to the tank, the following interaction was recorded for one hour and aggressive behaviors performed by the resident female were quantified.

### Inter-behavioral Intervals

We also calculated Inter-behavioral intervals as described in ^29^. The log files containing the results from BORIS were analyzed for inter-behavior intervals (IBI) using custom R software (available at https://github.com/AlwardLab). IBI were determined by calculating the time between successive behaviors and averaging across all IBIs within a given log file. Negative control fish performed almost no social behavior so were not included in analyses of temporal patterns of behavior.

### Morphological and steroid hormone analysis

Subjects were assessed for standard length (SL), body mass (BM), gonad mass, and gonadosomatic index [GSI = (gonad mass / body mass) * 100]. Male GSI ranged from 0.324 - 1.813 (average GSI = 0.895) and female GSI ranged from 1.074 - 9.122 (average GSI = 5.903). Blood samples were collected with capillary tubes from the caudal vein, centrifuged for 10 min at 8 rpm, and the plasma was removed and stored at -80°C until assayed.

Plasma 11-ketotestosterone (11-KT) levels were measured using a commercially available enzyme immunoassay (EIA) kit (Cayman Chemical Company, Ann Arbor, MI, USA). This kit has been extensively validated for use in *A. burtoni* ^30^. For the 11-KT assay, a 2-μl sample of plasma from each subject was extracted two times using 200 μl of ethyl ether and evaporated under a fume hood before re-constitution in EIA assay buffer (1:40 dilution). EIA kit protocols were then strictly followed, plates were read at 405 nm using a microplate reader (Biorad), and steroid concentrations were determined based on standard curves. All samples were assayed in duplicate. Intra and inter-assay variability were 13.5% and 10.9%, respectively.

### Immunohistochemistry for pS6

Following the cervical transection of subjects, brains were fixed in 4% paraformaldehyde in PBS (pH =7.4) for 1-2 hours and then transferred to 30% sucrose in PBS. Brains were kept in the sucrose solution at 4°C until they sunk and then embedded in mounting media Neg50 and stored at -80°C. Brains were sectioned at 30 μm. We analyzed neural activation in the brain following an aggression assay vs. a negative control condition using an immunohistochemistry protocol to detect the phosphorylated ribosome marker pS6, similar to that described in Butler et al., 2018.

We performed immunohistochemistry to detect ps6, a proxy of neural activation in the brain ^33^ that has been used previously in *A. burtoni* brains ^32,34^ After sectioning, the slides were dried in a desiccator for 48 hours, then stored at -80°C. Slides were then dried again in the desiccator for 48 hours prior to the immunohistochemistry protocol. Slides were outlined with a hydrophobic barrier using a PAP pen and then immersed in boiling citric acid (10 mM, pH = 6) for 5 minutes twice. The slides were then rinsed with 1x PBS (pH = 7.4) for 5 minutes three times. Nonspecific binding was blocked by incubating slides in 1x PBS containing 5% donkey serum and 0.3% Triton X-100 (PBS-T) for 30 minutes at room temperature. Then slides were incubated in PBS-T containing 0.5% donkey serum with the pS6 primary antibody (1:300, Invitrogen™ Phospho-S6 (Ser244, Ser247) polyclonal antibody) overnight at room temperature. Following the primary antibody incubation, slides were rinsed with 1x PBS for 5 minutes three times. Slides were then incubated in PBS-T containing 0.5% donkey serum with the donkey anti-rabbit secondary antibody (1:300, Invitrogen Alexa Fluor™ 488) at room temperature for 1 hour. The slides were rinsed with DAPI (1:500, Sigma-Aldrich) diluted in PBS for 5 minutes and then coverslipped with aqua-poly/mount and stored at 4°C.

### Quantification of ps6 immunoreactivity

Stained slides for the pS6 immunohistochemistry were imaged using a Nikon Eclipse 80i Microscope MicroFire™ at 20x magnification in GFP and DAPI and quantified in regions of the social behavior network (SBN). The social behavior network consists of interconnected nuclei that are implicated in the production of social behavior ^27,28^. We quantified pS6 immunoreactivity in the following brain regions of the SBN: the ventral part of the ventral telencephalon (Vv) (lateral septum homolog), the supracomissural part of the ventral pallium (Vs) (medial amygdala and bed nucleus of the stria terminalis partial homolog), subpopulations of the preoptic area (POA): parvocellular preoptic nucleus, anterior part (nPPa), magnocellular preoptic nucleus, magnocellular division (nMMp), magnocellular preoptic nucleus, parvocellular division (nPMp); anterior tuberal nucleus (ATn) (ventromedial hypothalamus homolog), and the periaqueductal gray / central gray (PAG/CG). We additionally quantified pS6 immunoreactivity in the vagal lobe (VL), vagal motor nucleus (Xm), medial part of the dorsal telencephalon, subdivision 3 (Dm-3), and the lateral nucleus of the dorsal telencephalon, granular region (Dl-g) due to the involvement of these regions in either processing sensory information or the social decision-making network, which is thought to evaluate the salience of social cues ^28^.

The number of pS6 positive cells were quantified for each ROI using ImageJ. A polygon was drawn around the ROI in each hemisphere and each pS6 positive cell within the ROI was counted manually. The cell density for each ROI was calculated per hemisphere (cells/μm^2^) and then the cell density was summed between the hemispheres. This total cell density measure was averaged across two consecutive sections for each region per subject, and the consecutive sections were selected at random using a random number generator. If the subject did not have two consecutive sections for a ROI, the cell density was taken from one section at random.

### Statistical Analysis

All statistical tests were performed using Graphpad Prism version 10. Two-way ANOVAs were used to compare 1) male and female aggressive behaviors over time in the mirror assay and 2) effects of sex and mirror condition on behaviors and pS6 immunoreactivity between negative control and mirror males and females. Following a significant main effect for ANOVA, Šídák’s test was used for multiple comparisons. When normality and equality of variance assumptions were not met, a log transformation of the data was performed before conducting the two-way ANOVA. If log transformations did not correct the data to meet assumptions, we still conducted a two-way ANOVA on the log-transformed data for consistency; this only occurred for two dependent variables (activity and ps6 immunoreactivity in the ATn). We also performed Fisher’s tests on the frequency of performance of rostral displays and quivers. Effects were considered significant at *p*≤0.05. For several behaviors, log transformations were not feasible due to zeros in the negative control groups. For these behaviors, two-way ANOVAs were conducted, and additional chi-square tests were run. Pearson correlations were used to test for correlations between pS6 immunoreactivity of each ROI for mirror and negative control fish. Missing values were interpolated across all conditions. A principal components analysis (PCA) on all ps6+ cell count data was performed for mirror and negative control conditions. Individual principal component scores were obtained by projecting a new vector onto the PCA space and then correlated to aggressive behaviors in mirror males and females using Spearman correlations.

## Results

### Male and female A. burtoni perform the same and different aggressive behaviors towards a mirror

Contour maps of fish movement over time that were generated in Noldus showed that before the opaque cover was lifted fish swim around the edges of the assay arena (Fig. 3a). However, fish spent the vast majority of their time in front of the mirror once the cover was removed.

**Figure 3.**
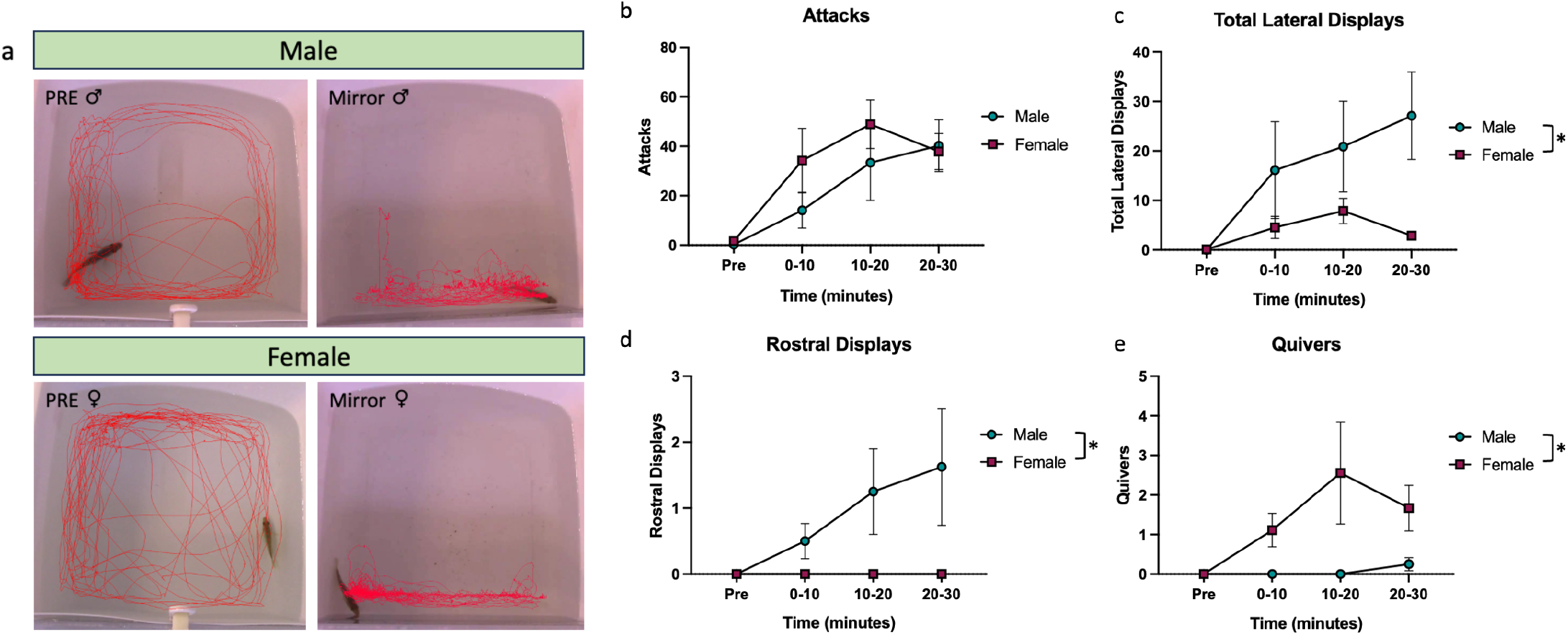
Male and female *A. burtoni* perform different aggressive behaviors. (a) Representative track visualization of male and female subjects before (PRE) and throughout (Mirror) a mirror assay trial. (b) Males and females both perform attacks in a mirror assay. (c)(d) Males perform more lateral displays and rostral displays than females. (e) Females perform more quivers than males.

**Figure 4.**
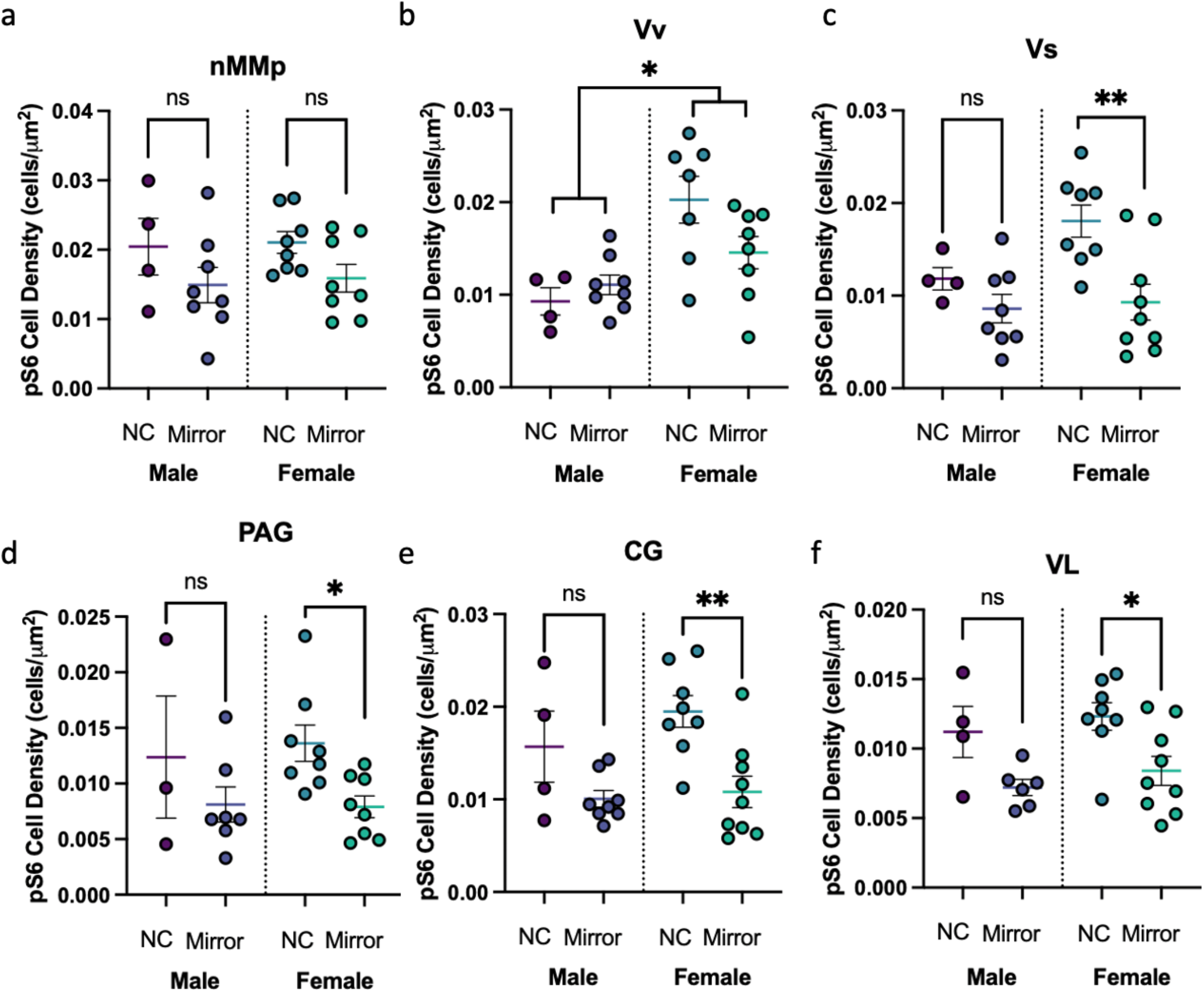
Sex differences and similarities in neural activation patterns. (a) pS6 immunoreactivity differed by condition in the nMMp, but post-hoc comparisons did not reveal significant differences in pS6 cell density within each sex. (b) Females have higher pS6 immunoreactivity in the Vv than males. (c)(d)(e)(f) pS6 cell density is higher in negative control fish than in mirror fish in the Vs, PAG, CG, and the VL. Multiple comparisons reveal that pS6 cell density is higher in negative control females compared to mirror females for these regions. NC, negative control. ** p < 0.01, * p < 0.05. ns, not statistically significant.

**Figure 5.**
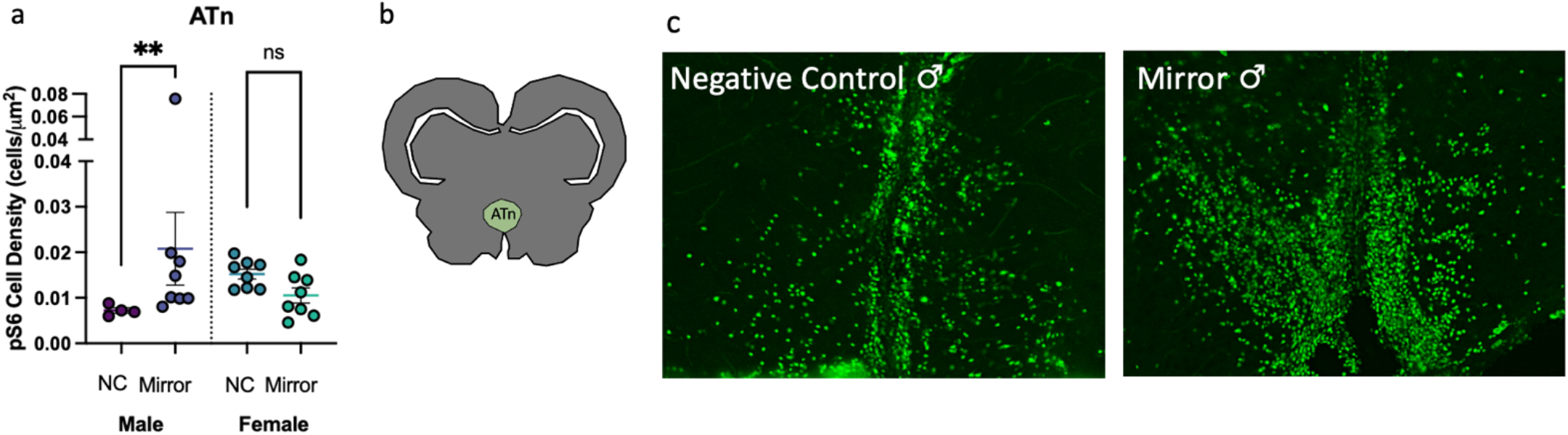
Males and females have opposite neural activation patterns in the ATn. (a) pS6 cell density in the ATn has an interaction effect between sex and condition. (b) Representative section of the ATn. (c) Representative sections of the ATn stained with pS6 antibody to detect neural activation show lower signal in a male in the control condition compared to a male exposed to a mirror. NC, negative control. ** p < 0.01

There was a significant effect of time but not sex or an interaction on attacks (Table 2; attacks, Two-way ANOVA, time: p < 0.0001). There was a significant effect of sex and time but not an interaction on lateral displays (Table 2; lateral displays, Two-way ANOVA, sex: p = 0.0399, time: p = 0.0126), wherein males performed more lateral displays than females (See videos S3-S4). We also observed a significant effect of sex on rostral displays (Table 2; rostral displays, Two-way ANOVA, sex: p = 0.0075), where males produced more rostral displays than females (See video S2). Additionally, there was a significant effect of sex on quivers (Table 2; quivers, Two-way ANOVA, sex: p = 0.0413). Specifically, females performed more quivers than males (See example in video S1). Fisher’s exact tests showed more males than females performed rostral displays (6/8 males versus 1/9 females; *p =* 0.0152) while more females than males performed quivers (2/8 males versus 8/9 females; *p =* 0.0152). We were surprised to observe females perform a male-typical mating behavior, quivers, in the mirror assay, given the behavior is putatively used as an aggressive display in a mirror assay by females. To confirm females do these behaviors towards another female, we conducted resident intruder tests with focal resident females against intruder females ^25^. Female residents performed quivers, lateral displays, and attacks towards intruder females and also chased them (Fig. S1), indicating quivers are a normal component of the behavioral repertoire of female-female aggression. Across 5 resident females, attacks ranged from 2-56 (Average: 22.6, St. Dev.:22.15), lateral displays ranged from 1-30 (Average: 12.8, St. Dev.: 10.76), and quivers ranged from 0-9 quivers (Average: 3.2, St. Dev.: 3.77).

**Table 2.** Effect of sex and time on aggressive behaviors quantified in a mirror assay.

|  | Sex |  | Time |  | Sex*Time |  |
| --- | --- | --- | --- | --- | --- | --- |
|  | F | P | F | P | F | P |
| Attacks | 0.7888<br>1, 15 | 0.3885 | 12.60<br>2.435, 36.53 | **< 0.0001** | 1.076<br>3, 45 | 0.3688 |
| Lateral displays | 5.059<br>1, 15 | **0.0399** | 4.618<br>2.353, 35.29 | **0.0126** | 2.397<br>3, 45 | 0.0805 |
| Rostral displays | 9.537<br>1, 15 | **0.0075** | 1.946<br>1.529, 22.93 | 0.1724 | 1.946<br>3, 45 | 0.1357 |
| Quivers | 4.982<br>1, 15 | **0.0413** | 3.038<br>1.306, 19.59 | 0.0882 | 2.760<br>3, 45 | 0.0530 |

There were no sex differences in median or mean IBI between mirror males and females (Fig. S2a; Median IBI, Unpaired t-test, *p* = 0.1724, *t* = 1.433, df = 15, Fig. S2b; Mean IBI, Mann-Whitney, *p* = 0.0927, U = 18).

We compared the behaviors of the NC fish to mirror fish and found across all behaviors NC males and females performed no or very few acts of any behavior types (Fig. S3). Overall, Two-way ANOVAs revealed consistent effects of condition and not sex or an interaction on behaviors performed during the mirror assay (See Supplementary Results for detailed statistical findings; Table S1). This shows the NC fish included in this assay are sufficient as a control group when comparing ps6 immunoreactivity to fish exposed to the mirror.

There was a significant effect of condition but not sex or an interaction on latency to behave (Table 3; latency to behave, Two-way ANOVA, condition: *p* = 0.0408), wherein mirror males had a longer latency to behave than negative control males (Table 3; latency to behave, Šídák’s test, Male Mirror – NC, *p* = 0.0259). There were no significant effects of sex, condition, or sex*condition on latency to aggression (Table 3; latency to aggression, Two-way ANOVA). Additionally, there were no significant effects of sex, condition, or sex*condition on distance moved and velocity (Table 3; distance moved, Two-way ANOVA of log transformation; velocity, Two-way ANOVA of log transformation).

**Table 3.** (1) Effect of sex and condition on tracking and activity data quantified in a mirror assay. (2) Post-hoc comparisons.

|  | Sex |  | Condition |  | Sex*Condition |  |
| --- | --- | --- | --- | --- | --- | --- |
|  | F | P | F | P | F | P |
| Latency to behave | 1.616<br>1, 26 | 0.2149 | 4.632<br>1, 26 | **0.0408** | 3.630<br>1, 26 | 0.0679 |
| Latency to aggression | 0.02848<br>1, 17 | 0.8680 | 0.1388<br>1, 17 | 0.7141 | 2.735<br>1, 17 | 0.1165 |
| Distance moved | 0.03466<br>1, 26 | 0.8537 | 0.06752<br>1, 26 | 0.7970 | 0.06312<br>1, 26 | 0.8036 |
| Velocity | 0.06927<br>1, 26 | 0.7945 | 0.005043<br>1, 26 | 0.9439 | 0.1013<br>1, 26 | 0.7528 |
| Activity | 1.242<br>1, 25 | 0.2757 | 13.13<br>1, 25 | **0.0013** | 0.4020<br>1, 25 | 0.5318 |
| Zone frequency | 0.1061<br>1, 26 | 0.7472 | 0.04381<br>1, 26 | 0.8358 | 0.7549<br>1, 26 | 0.3929 |
| Zone latency | 2.378<br>1, 26 | 0.1351 | 2.121<br>1, 26 | 0.1573 | 2.256<br>1, 26 | 0.1452 |
| Zone duration | 3.086<br>1, 26 | 0.0907 | 4.841<br>1, 26 | **0.0369** | 1.451<br>1, 26 | 0.2392 |
| Mean distance to the zone | 1.701<br>1, 26 | 0.2036 | 10.53<br>1, 26 | **0.0032** | 0.08735<br>1, 26 | 0.7699 |
| Total distance to the zone | 1.506<br>1, 26 | 0.2308 | 10.71<br>1, 26 | **0.0030** | 0.1012<br>1, 26 | 0.7529 |

|  | Male Mirror - NC | Female Mirror - NC |
| --- | --- | --- |
|  | P | P |
| Latency to behave | **0.0259** | 0.9776 |
| Activity | 0.1109 | **0.0072** |
| Zone duration | 0.7684 | **0.0285** |
| Mean distance to the zone | 0.1230 | **0.0223** |
| Total distance to the zone | 0.1224 | **0.0204** |

We observed a significant effect of condition but not sex or an interaction on activity (Table 3; activity, Two-way ANOVA of log transformation, condition: *p* = 0.0013). One mirror female outlier was removed. Mirror females have higher activity than negative control females and males follow this same pattern although not significant (Table 3, activity, Šídák’s test, Female Mirror – NC, *p* = 0.0072). No significant effects of sex, condition or sex*condition were observed on zone frequency or zone latency (Table 3; zone frequency, Two-way ANOVA of log transformation; zone latency, Two-way ANOVA). There was a significant effect of condition but not sex or an interaction on zone duration (Table 3; zone duration, Two-way ANOVA, condition: *p* = 0.0369) and mirror females had a longer zone duration than negative control females while males did not differ by condition (Table 3, zone duration, Šídák’s test, Female Mirror – NC, *p* = 0.0285).

Condition—but not sex or an interaction—had an effect on mean distance and total distance to the zone (Table 3; mean distance to the zone, Two-way ANOVA of log transformation, condition: *p* = 0.0032; total distance to the zone, Two-way ANOVA of log transformation, condition: *p* = 0.0030). Mirror females had a higher mean and total distance to the zone than negative control females and males followed this pattern but was not statistically significant (Table 3, mean distance to the zone: *p* = 0.0223, total distance to the zone: *p* = 0.0204, Šídák’s test).

### Sex differences and similarities in neural activation patterns of the SBN after aggression

No significant effects of sex, condition, or sex*condition on ps6 immunoreactivity were observed in the POA or subregions nPPa and nPMp, as well as in the Dl-g, Dm-3, and Xm (Table S2; POA, Two-way ANOVA; nPPa Two-way ANOVA; nPMp, Two-way ANOVA; Dl-g, Two-way ANOVA; Dm-3, Two-way ANOVA; Xm, Two-way ANOVA of log transformation). However, there was a significant effect of condition, but not sex or an interaction, on pS6 immunoreactivity in one subregion of the POA, the nMMp (Table 4; nMMp, Two-way ANOVA, condition: *p* = 0.0400) in which negative controls had higher ps6 immunoreactivity in this region on average, but this was not statistically significant. There was a significant effect of sex on ps6 immunoreactivity in the Vv, where females had higher pS6 immunoreactivity compared to males (Table 4; Vv, Two-way ANOVA, sex: *p* = 0.0010).

**Table 4.** (1) Effect of sex and condition on pS6 immunoreactivity and 11-KT. (2) Post-hoc comparisons.

| 1 | Sex |  | Condition |  | Sex*Condition |  |
| --- | --- | --- | --- | --- | --- | --- |
|  | F | P | F | P | F | P |
| nMMp | 0.1046<br>1, 24 | 0.7492 | 4.718<br>1, 24 | **0.0400** | 0.005974<br>1, 24 | 0.9390 |
| Vv | 14.21<br>1, 23 | **0.0010** | 1.043<br>1, 23 | 0.3178 | 3.834<br>1, 23 | 0.0624 |
| Vs | 3.347<br>1, 25 | 0.0793 | 10.03<br>1, 25 | **0.0040** | 2.127<br>1, 25 | 0.1571 |
| PAG | 0.5695<br>1, 22 | 0.4585 | 5.272<br>1, 22 | **0.0316** | 0.3985<br>1, 22 | 0.5344 |
| CG | 1.412<br>1, 25 | 0.2459 | 14.06<br>1, 25 | **0.0009** | 0.6412<br>1, 25 | 0.4308 |
| VL | 1.026<br>1, 23 | 0.3216 | 11.96<br>1, 23 | **0.0021** | 0.0009858<br>1, 23 | 0.9752 |
| ATn | 0.4562<br>1, 24 | 0.5059 | 0.7065<br>1, 24 | 0.4089 | 10.14<br>1, 24 | **0.0040** |
| 11-KT | 0.2400<br>1, 26 | 0.6283 | 1.826<br>1, 26 | 0.1882 | 0.01443<br>1, 26 | 0.9053 |

| 2 | Male Mirror – NC | Female Mirror - NC |
| --- | --- | --- |
|  | P | P |
| nMMp | 0.2935 | 0.2092 |
| Vv | 0.7935 | 0.0598 |
| Vs | 0.4903 | **0.0020** |
| PAG | 0.5310 | **0.0408** |
| CG | 0.1378 | **0.0024** |
| VL | 0.0773 | **0.0181** |
| ATn | **0.0313** | 0.1467 |

We observed a significant effect of condition on ps6+ cells in the Vs, PAG/CG, and the VL (Table 4; Vs, Two-way ANOVA, condition: *p* = 0.0040; PAG, Two-way ANOVA of log transformation, condition: *p* = 0.0316; CG, Two-way ANOVA, condition: *p* = 0.0009; VL, Two-way ANOVA, condition: *p* = 0.0021).

Specifically, in the Vs, PAG/CG, and the VL, female negative controls had higher pS6 immunoreactivity than mirror females while males did not differ by condition (Table 4, Vs: *p* = 0.0020, PAG: *p* = 0.0408, CG: *p* = 0.0024, VL: *p* = 0.0181, Šídák’s tests). Mirror males and negative controls did not differ in ps6 immunoreactivity in the Vs, PAG, and CG (Table 4, Vs, PAG, CG, Šídák’s tests). Although not statistically significant, male negative controls had higher pS6 immunoreactivity in the VL compared to mirror males (Table 4, VL, Šídák’s test, Male Mirror – NC, *p* = 0.0773), following the same trend as females in the VL.

A significant sex*condition interaction on ps6 immunoreactivity in the ATn was observed (Table 4; ATn, Two-way ANOVA of log transformation, interaction: *p* = 0.0040). Specifically, males exposed to the mirror had more ps6+ cells in the ATn compared to negative control males, while this was not true for females (Table 4, ATn, Šídák’s test, Male Mirror – NC, *p* = 0.0313).

### No effects of sex or condition on 11KT

There were no significant effects of sex, condition or sex*condition on 11-KT (Table 4; 11-KT, Two-way ANOVA).

### Correlated neural activity patterns differ in mirror-exposed and negative control fish

We correlated the pS6 immunoreactivity across regions of the SBN in mirror and negative control fish to investigate the functional connectivity of these regions in response to an aggressive assay. Principal components analysis (PCA) of neural activation patterns revealed two significant components that capture the variability in the data (Figure 6). For the mirror fish, the first component (PC1, 35.4% explained variance) was primarily weighted by ps6+ cell count in the subregions of the POA (nPPa, nMMp, and nPMp) and the Dm-3 while the second component (PC2, 16.9% explained variance) was weighted primarily by ps6+ cell count in the CG, VL, Dl-g, and the Xm and ATn. Variation in male neural activity is mostly captured across PC1, while female neural activation patterns are explained relatively equally across both components. In the negative control fish, the first principal component (59.3% explained variance) is weighted by the Vs, POA (and subregion nPMp), and the Dm-3. The second component is driven by neural activation of the Vv, ATn, CG, VL, and Xm. Males and females are largely overlapping based on the biplot, suggesting that there are not sex differences in the neural activation patterns captured by PC1 and PC2 in the negative control fish.

**Figure 6.**
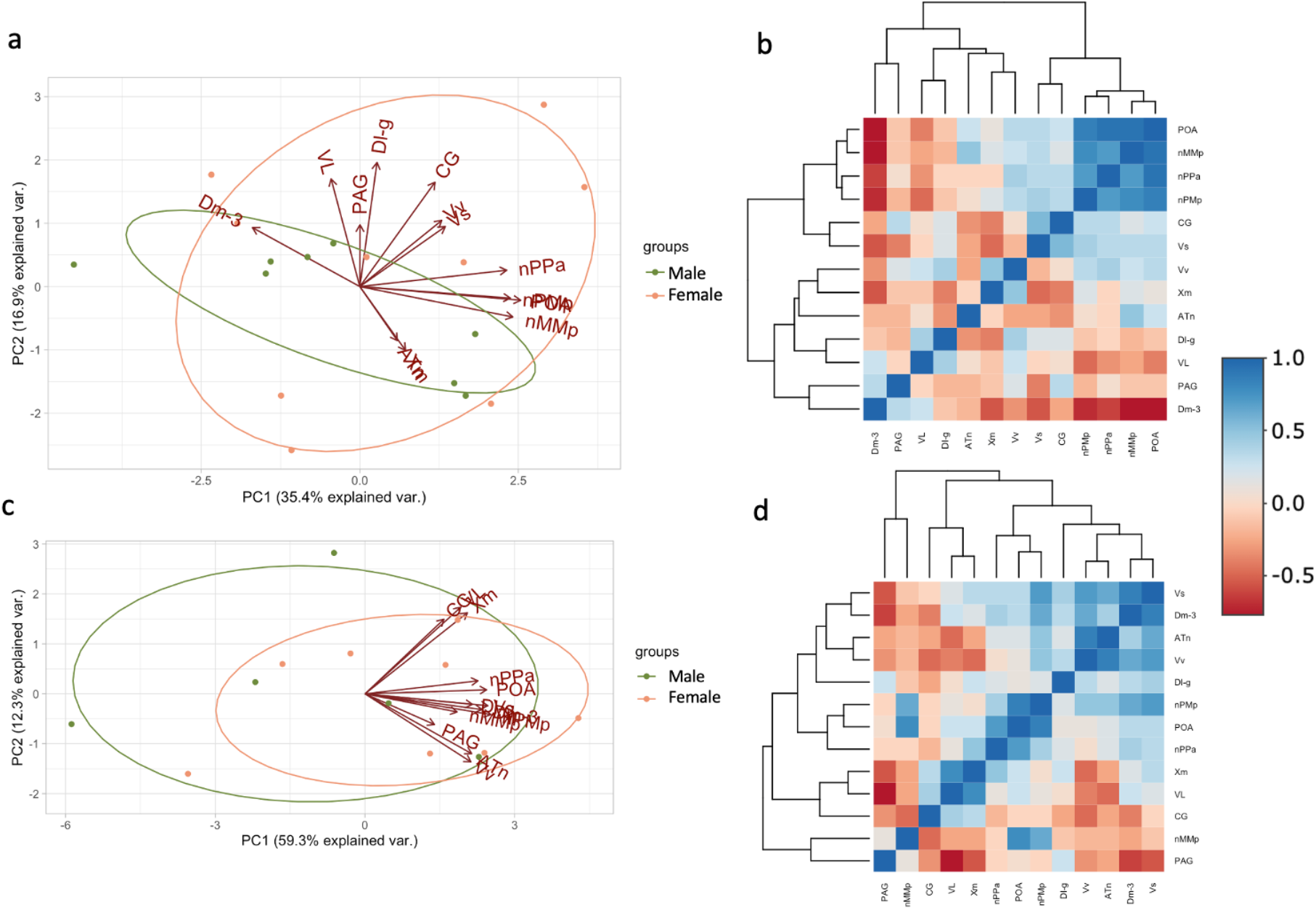
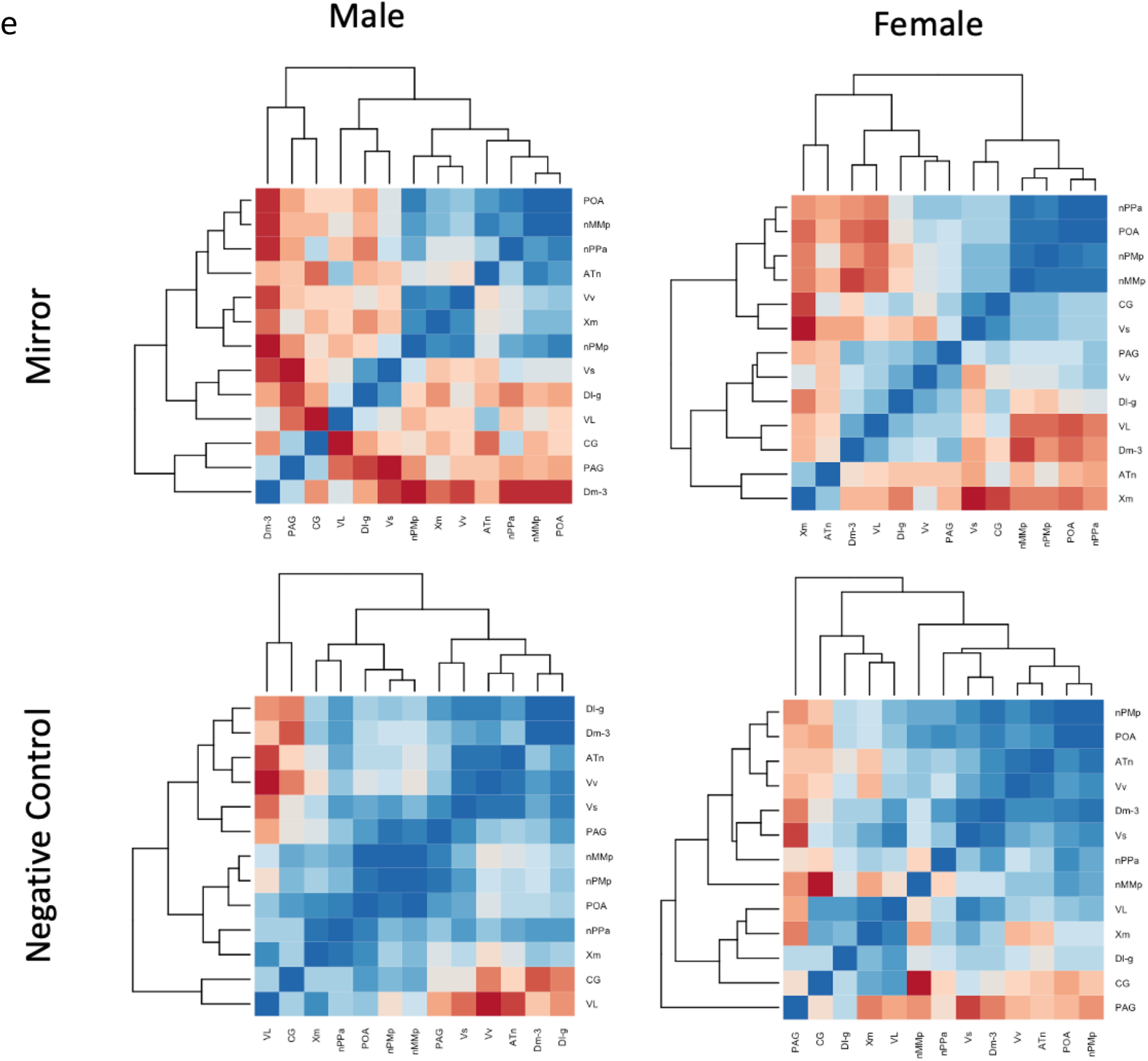
Neural activation clusters by condition and sex. (a) Principal components analysis of pS6 immunoreactivity in mirror fish. (b) Heat map of Pearson correlation coefficients of pS6 immunoreactivity in the SBN of mirror fish. (c) Principal components analysis of pS6 immunoreactivity in negative control fish. (d) Heat map of Pearson correlation coefficients of pS6 immunoreactivity in the SBN of negative control fish. (e) Heat maps of Pearson correlation coefficients of pS6 immunoreactivity in the SBN for each sex and condition.

To further investigate the correlated pS6 immunoreactivity across the regions of interest, we used Pearson correlations and created heatmaps from the correlation coefficients for mirror and negative control fish with dendrograms to visualize the clustering of neural activity (Figure 6). For the mirror fish, there is a clear cluster of the POA and subregions (nPPa, nMMp, and nPMp) with a strong negative correlation to the Dm-3. For the negative control fish, there is a cluster of correlated neural activity between the Vs, Dm-3, ATn, and the Vv, and an additional cluster between the PAG/CG, nMMp, VL, and Xm. We then wondered if these clusters were consistent between males and females within each condition (Figure 6e). In the mirror males, subregions of the POA (nPPa and nMMp) clustered with the ATn. In the mirror females, there is still a cluster of the POA and subregions (nPPa, nMMp, and nPMp), but the ATn is negatively correlated and clusters with the Dm-3, VL, and Xm. The negative control males have correlated activity in the Vs, Vv, ATn, Dl-g, and Dm-3, that is negative correlated to the CG and VL. The female negative controls have a cluster of the POA (nPMp), ATn, and Vv.

With the differences in correlated neural activity between mirror males and females specifically in their opposite relationship to the ATn, we wanted to further investigate if principal component scores of the mirror fish (Figure 6a) correlated to aggressive behaviors (Figure 7). We correlated the principal components scores of each mirror fish for PC1 and PC2 to aggressive behaviors and found significant correlations between PC2 with attacks and total aggression in mirror males and females. However, males and females had opposite relationships between these measures - males had a positive correlation between PC2 and attacks/total aggression while females had a negative correlation between PC2 and attacks/total aggression.

**Figure 7.**
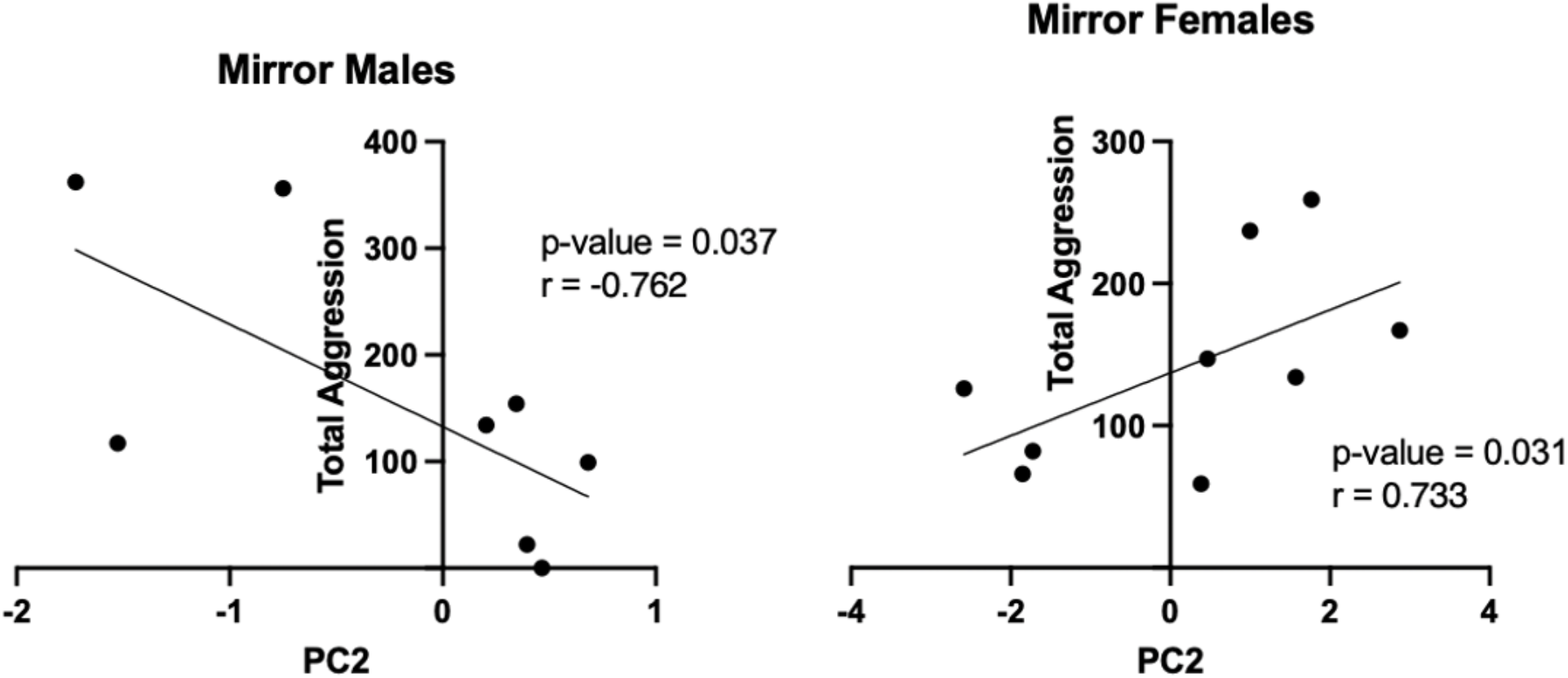
Correlations of PC2 to total aggression in mirror males and mirror females.

## Discussion

We aimed to compare male and female intrasexual aggression in *A. burtoni*, a non-traditional model organism of social behavior that has the potential to yield novel insights into the sexually dimorphic control of social behavior. We determined that in an identical context that elicits aggression, male and female *A. burtoni* perform a mixture of identical behaviors and sex-specific behaviors. Interestingly, quivers, which are performed by males in reproductive contexts, were performed by females in the mirror assay, suggesting it is used as an aggressive signal by females. As in other species in which females adopt male-like mounting behaviors towards other females to assert dominance, perhaps in teleosts this behavior maintains female social status ^35^.

Intrasexual male aggression is thought to follow specific “rules of engagement” between opponents, assessing the resource holding power of the other to minimize risk in conflict ^36^. Since *A. burtoni* males alter their aggression depending on the relative size of their opponent, we can safely assume that intrasexual male aggression possesses a level of opponent assessment. In our study, we find that males exposed to a mirror have a longer latency to behave compared to negative control males. Perhaps this longer duration of time to investigate the mirror serves as a lengthy assessment period since the male is faced with an identically-sized opponent. However, we don’t see this difference in female *A. burtoni* exposed to a mirror vs. negative controls. Perhaps intrasexual female aggression in *A. burtoni* does not follow the same opponent assessment as it does for *A. burtoni* males, supported by a study in Texas cichlids that suggests that males and females use a different set of rules to settle conflict ^37^.

We also revealed a significant difference in condition in activity and zone duration in which mirror females have higher activity and spend more time in the zone near the mirror compared to negative control females. Although not statistically significant, mirror males also have higher activity levels on average than negative controls. Since the change in the environment between the pre-mirror and mirror assay is more pronounced in trials of fish exposed to a mirror compared to the negative controls (opaque cover removed vs. remained on), we expect to see mirror subjects behaving more actively in their assay. Additionally, the mirror allows subjects to be more engaging with their reflection compared to an opaque cover which may be manifesting here in activity levels. Mirror females spending more time near the mirror may reflect an aspect of exploration or investigation that is not necessary for males in this context. In fact, female rats show more exploratory activity than males in a novel environment and have shorter approach latencies to novelty ^38^. Perhaps this aspect of elevated exploration in females is consistent with a longer zone duration in mirror females that we have shown here.

We found that 11-KT did not differ between mirror and negative control conditions, which is in line with findings in other teleost species ^39,40^ but contrasts with previous findings in male *A. burtoni* where mirror assays were also used ^24^. However, as argued by Oliveira and Canario ^41^, the previous mirror assay study in male *A. burtoni* ^24^ has a few issues that could explain the higher 11-KT in mirror fish compared to negative controls, including the confound of exposure to a competing male’s chemical cues in the same territory directly before exposure to a mirror. Indeed, the lack of a rise in androgens in a mirror assay that we observed in both males and females has been explained by the fact that there is no clear winner of the fight as is seen against a live opponent because all aggressive acts are perceived to elicit an equal counter response ^14,39–42^.

In multiple brain regions we observed impacts of sex and exposure to the mirror on neural activation as measured by ps6 immunoreactivity. For several brain regions no differences in pS6 activation between negative control and mirror fish, including the POA (nPPa and nPMp), Dl-g, Dm-3, and Xm, were observed. The POA plays a role in aggressive and reproductive behaviors across vertebrates ^28^. In *A. burtoni*, there is an increase in *egr1* expression in the nMMp in males after fighting but not after courting, emphasizing the role of this POA subregion in modulating aggression ^43^. When we investigated the pS6 immunoreactivity in the POA we expected to see higher pS6 immunoreactivity in the POA in fish exposed to the mirror compared to the negative control condition. Surprisingly, we see no differences in pS6 immunoreactivity as a function of sex or condition for two of the subregions of the POA (nPPa and nPMp), and although we find a significant effect of condition on pS6 immunoreactivity in the nMMp, it was higher in negative controls. Another study in *A. burtoni* found no effect of social stimulus on c-Fos expression in the POA when males were exposed to reproductive or aggressive contexts ^44^. Interestingly, the males in the aforementioned study were not engaged in full contact with stimulus fish and therefore not receiving all sensory signals. Hence, it may be that the POA neurons activate in response to integrating several multisensory components of social information, but solely the visual input and behavioral output we have observed in this study is not sufficient for the activation of these neurons. Previous work has emphasized the importance of the integration of chemosensory and visual signals to elicit distinct neural activation patterns compared to the exposure of a unimodal signal ^44,45^. Since our design utilizes a unimodal visual signal, we would hypothesize that neural activation of the POA would differ by condition if fish were exposed to multisensory social cues.

The Dl-g (homolog of the medial pallium; hippocampus) is involved in processing sensory input and receiving and processing socially relevant hydrodynamic cues in *A. burtoni* ^46^. The Dm (homologous to the basolateral amygdala, blAMY) also receives sensory input including lateral line information ^47–49^. pS6 immunoreactivity in the Dl-g, Dm-3, and Xm did not differ by sex or condition and may indicate that mechanosensory input is similar in the negative control condition and mirror conditions despite distinct differences in behavioral output. Some behavioral measures (distance moved (cm) and velocity (cm/s)) do not differ by sex or condition and may provide evidence of similar mechanosensory input between these groups. It is important to note that the lack of socially relevant hydrodynamic cues that accompany a fight with a real opponent are not present in the mirror assay, and thus regions like the Dl-g, Dm-3, and Xm that process sensory information may not be relevant in an aggressive assay lacking this dynamic sensory input.

One region that did display a sex difference in pS6 immunoreactivity is the Vv. The Vv, homologous to the LS, is thought to integrate internal state with the social environment since this region is shared between the social behavior network and the mesolimbic reward system ^28^. The Vv has also been implicated in processing social information in anxiety-like contexts. In previous work, female *A. burtoni* had higher *cfos* expression in the Vv after watching a preferred male lose a fight ^50^. In this study, we found higher pS6 immunoreactivity in the Vv of females compared to males regardless of condition, which may indicate that the mirror assay is more anxiogenic to females, or that the Vv is more active overall in females than it is in males. Future experiments aimed at testing the role of the Vv in explicitly anxiogenic behavior paradigms in both males and females are thus warranted.

The ventromedial hypothalamus (VMH) has been implicated in mediating aggressive behaviors across several species ^6^. In mammals, *cfos* expression in the VMH increases after aggressive interactions in males, and specifically the causal role of the ventrolateral portion of the VMH (VMHvl) in aggression in male mice is well established ^3,51–53^. In *A. burtoni*, the ATn (homologous to the VMH) is likely involved in processing mechanosensory information in male-male aggressive contexts ^46^. In our study, pS6 immunoreactivity in the ATn is higher in males exposed to a mirror than in negative controls, supporting the importance of activity in the ATn in mediating male-male aggression. While ps6 immunoreactivity was not higher in mirror versus control females, this does not preclude the ATn potentially being involved in female aggression in an environment-specific manner. For example, ATn neurons may become activated during female aggression in particular contexts and/or a specific sub-population of neurons in the ATn that remains to be identified regulates female aggression as the ventrolateral portion of the VMH does in male mice. Investigations into whether a specific locus within the fish VMH regulates aggression in both male and female *A. burtoni* will be key in disentangling the neural basis of aggression in this species.

Some regions of interest had higher neural activation in negative controls than in mirror fish. These results suggest the potential involvement of inhibitory neurons in suppressing activity in specific aggression circuits. Glutamic acid decarboxylase (Gad) is the rate-limiting enzyme that converts glutamate (an excitatory neurotransmitter) into GABA (an inhibitory neurotransmitter). Gad exists in two highly conserved isoforms, *gad1* and *gad2*. In *A. burtoni*, the Vs, CG, and VL express both *gad1* and *gad2* ^54^. The Vs, PAG/CG and VL show higher pS6 immunoreactivity in negative controls and since *gad1* and *gad2* are both expressed in these regions, we postulate that the inhibitory neurons in these regions are important for suppressing aggressive behaviors. We hypothesize that inhibitory neurons containing GABA are activated in negative control fish whereas lower activation of these neurons permits aggressive behaviors in mirror fish.

The clusters of correlated neural activity in the mirror fish replicated the loadings in the PCA – in which the POA and subregions cluster together and are negatively correlated to the Dm-3. When we visualized the clusters of correlated neural activity separately in mirror males and females, we found a distinct difference in the relationship of the ATn to the POA present in males and not females (Figure 6e). This strong correlation of the POA and the ATn present in males and not females reflects a male-specific aggressive state. The core aggression circuit (CAC) is a subnetwork of the SBN that is specialized for aggression and is comprised of the medial amygdala, the ventrolateral portion of the ventromedial hypothalamus (VMHvl), the ventral premammilary nucleus, and the bed nucleus of the stria terminalis ^3^. The POA acts as a link between the CAC and the ventral tegmental area to trigger learned aggressive actions indirectly. Since the ATn and the POA are interconnected, perhaps in male *A. burtoni* aggressive behaviors are initiated by the ATn (VMH homolog) and the POA, and this is not the case in females. The functional connectivity between the POA and ATn in aggressive males may suggest the presence of a permissive gate that is triggered by the visual stimulation of an aggressive encounter that opens this connection between the ATn and POA. Since this functional connectivity is exclusive to aggressive males, a difference mechanism may be triggering aggression in females.

We also found that PC2 for mirror fish correlated positively to attacks and total aggression in mirror males and correlated negatively to attacks and total aggression in mirror females (Figure 7). This leads us to believe that the loading variables of PC2 (CG, VL, Xm, Dl-g, and ATn) mediating aggressive responses in males and females in opposite ways, perhaps through neural inhibition in females. These brain regions play a large role in the processing of sensorimotor information and could reflect sex differences in the circuits that function together to perform attacks in *A. burtoni*.

## Conclusions

Our results revealed both similar and sexually dimorphic behavioral patterns underlying aggressive responses to a mirror in male and female *A. burtoni*. We also observed extensive sex differences in neural activation throughout regions of the SBN in response to our aggression assay, suggesting distinct neurons and neural circuitry underlie aggressive behavior in male and female *A. burtoni*. Interestingly, neural activation in the ATn reflected aggressive state in males but not females, suggesting this region controls aggression in male *A. burtoni* as it does in mice ^3,7,52,55^. Activity in the ATn and the POA appears to be functionally connected in aggressive males, suggesting that the visual input received about an aggressive context in males triggers activity in the ATn and POA to produce a behavioral response, but this circuit is absent in female aggression. We also provide evidence that the inhibition of neural activity in a variety of brain regions in the SBN is associated with aggression in *A. burtoni* as well. These findings overall lay a foundation of testable hypotheses for future studies to understand the molecular and neural regulation of sexually dimorphic behaviors in *A. burtoni*.

Gaining a fundamental understanding of the molecular and neural basis of aggression requires studies in both sexes in which the conditions and behaviors performed by either sex are similar. In our mirror assay we were able to stimulate aggressive responses in both males and females. The behaviors performed by males were the same as the suite of behaviors they perform in dyadic social interactions and we confirmed that this is the case for females as well. A significant benefit of assessing aggression in a setup that controls for the environmental and social conditions in males and females is it is presumed that while 1) both sexes are experiencing an elevated motivation to behave aggressively, they are 2) also performing distinct and similar aggressive behavioral acts. Thus, a benefit for the use of *A. burtoni* as a model organism for elucidating the molecular and neural mechanisms governing sexually dimorphic aggression is that the molecular and neural mechanisms of the motivation versus sensorimotor aspects of aggression in both sexes can be studied in the same environment. Therefore, studying sex differences in aggression in *A. burtoni* may lead to new ideas on the fundamental mechanisms controlling social behavior generally, yielding novel hypotheses that can be tested across species.

## Supplementary Information

Supplementary results and figures are in the Supplementary Information file.

## Availability of data and materials

The datasets generated and/or analyzed during the current study are available in the manuscript itself, the Supplementary Information file, and online at https://github.com/AlwardLab/sexdiffs2024.

## Declarations

### Ethics approval and consent to participate

Experimental procedures were conducted according to the ethical guidelines for the care and use of laboratory animals. Experiments were approved by the University of Houston Institutional Animal Care and Use Committee (Protocol #202000001).

### Consent for publication

Not applicable

## Acknowledgements

The authors thank Andrew Hoadley for procedural assistance as well as Mariana Lopez for helpful discussions on interpreting findings.

## Authors’ contributions

B.A.A. and L.R.J. designed the study; L.R.J. performed the experiments; L.R.J. and M.D. collected the data; B.A.A. and L.R.J. analyzed the data; L.R.J. made the figures. B.A.A. and L.R.J. wrote and edited the manuscript.

## References

1. Adkins-Regan, E. Hormones and Animal Social Behavior. (Princeton University Press, 2005).

2. Nelson, R. & Kriegsfeld, L. J. An Introduction to Behavioral Endocrinology. (Sinauer, 2017).

3. Lischinsky, J. E. & Lin, D. Neural mechanisms of aggression across species. Nat. Neurosci. 23, (2020).

4. Bayless, D. W. & Shah, N. M. Genetic dissection of neural circuits underlying sexually dimorphic social behaviours. Philos. Trans. R. Soc. B Biol. Sci. 371, 1–10 (2016).

5. Been, L. E., Gibbons, A. B. & Meisel, R. L. Towards a neurobiology of female aggression. Neuropharmacology 156, 107451 (2019).

6. Nelson, R. J. & Trainor, B. C. Neural mechanisms of aggression. Nat. Rev. Neurosci. 8, 536–546 (2007).

7. Anderson, D. J. Circuit modules linking internal states and social behaviour in flies and mice. Nat. Rev. Neurosci. 17, 692–704 (2016).

8. McCarthy, M. M., Arnold, A. P., Ball, G. F., Blaustein, J. D. & De Vries, Geert. J. Sex Differences in the Brain: The Not So Inconvenient Truth. J. Neurosci. 32, 2241–2247 (2012).

9. Alward, B. A., Hoadley, A. P., Jackson, L. R. & Lopez, M. S. Genetic dissection of steroid-hormone modulated social behavior: Novel paralogous genes are a boon for discovery. Horm. Behav. 147, 105295 (2023).

10. Fernald, R. D. Social control of the brain. Annu. Rev. Neurosci. 35, 133–51 (2012).

11. Fernald, R. D. & Maruska, K. P. Social information changes the brain. Proc. Natl. Acad. Sci. 109, 17194– 17199 (2012).

12. Jackson, L. R., Lopez, M. S. & Alward, B. Breaking Through the Bottleneck: Krogh’s Principle in Behavioral Neuroendocrinology and the Potential of Gene Editing. Integr. Comp. Biol. 63, 428–443 (2023).

13. Maruska, K. P. & Fernald, R. D. Astatotilapia burtoni: A Model System for Analyzing the Neurobiology of Behavior. ACS Chem. Neurosci. 9, 1951–1962 (2018).

14. Munley, K. M. & Alward, B. A. Control of social status by sex steroids: insights from teleost fishes. Mol. Psychol. Brain Behav. Soc. 2, 21 (2023).

15. Maruska, K. P. & Fernald, R. D. Social Regulation of Male Reproductive Plasticity in an African Cichlid Fish. Integr. Comp. Biol. 53, 938–950 (2013).

16. Maruska, K. P. Social Transitions Cause Rapid Behavioral and Neuroendocrine Changes. Integr. Comp. Biol. 55, 1–13 (2015).

17. Maruska, K. P. & Butler, J. M. Reproductive- and Social-State Plasticity of Multiple Sensory Systems in a Cichlid Fish. Integr. Comp. Biol. 61, 249–268 (2021).

18. Renn, S. C. P., Fraser, E. J., Aubin-Horth, N., Trainor, B. C. & Hofmann, H. A. Females of an African cichlid fish display male-typical social dominance behavior and elevated androgens in the absence of males. Horm. Behav. 61, (2012).

19. Quintana, L. et al. Building the case for a novel teleost model of non-breeding aggression and its neuroendocrine control. J. Physiol.-Paris 110, 224–232 (2016).

20. Batista, G., Zubizarreta, L., Perrone, R. & Silva, A. Non-sex-biased Dominance in a Sexually Monomorphic Electric Fish: Fight Structure and Submissive Electric Signalling. Ethology 118, 398–410 (2012).

21. Karino, K. & Nakazono, A. Reproductive behavior of the territorial herbivoreStegastes nigricans (Pisces: Pomacentridae) in relation to colony formation. J. Ethol. 11, 99–110 (1993).

22. Vullioud, P., Bshary, R. & Ros, A. F. H. Intra- and interspecific aggression do not modulate androgen levels in dusky gregories, yet male aggression is reduced by an androgen blocker. Horm. Behav. 64, 430–438 (2013).

23. Ariyomo, T. O. & Watt, P. J. Aggression and sex differences in lateralization in the zebrafish. Anim. Behav. 86, 617–622 (2013).

24. Desjardins, J. K. & Fernald, R. D. What do fish make of mirror images? Biol. Lett. 6, (2010).

25. Alward, B. A., Cathers, P. H., Blakkan, D. M., Hoadley, A. P. & Fernald, R. D. A behavioral logic underlying aggression in an African cichlid fish. Ethology 127, 572–581 (2021).

26. Li, C.-Y., Hofmann, H. A., Harris, M. L. & Earley, R. L. Real or fake? Natural and artificial social stimuli elicit divergent behavioural and neural responses in mangrove rivulus, Kryptolebias marmoratus. Proc. R. Soc. B Biol. Sci. 285, 20181610 (2018).

27. Newman, S. W. The Medial Extended Amygdala in Male Reproductive Behavior A Node in the Mammalian Social Behavior Network. Ann. N. Y. Acad. Sci. 877, 242–257 (1999).

28. O’Connell, L. A. & Hofmann, H. A. The Vertebrate mesolimbic reward system and social behavior network: A comparative synthesis. J. Comp. Neurol. 519, (2011).

29. Alward, B. A., Hilliard, A. T., York, R. A. & Fernald, R. D. Hormonal regulation of social ascent and temporal patterns of behavior in an African cichlid. Horm. Behav. 107, 83–95 (2019).

30. Maruska, K. P. & Fernald, R. D. Steroid receptor expression in the fish inner earvaries with sex, social status, and reproductive state. BMC Neurosci. 11, (2010).

31. Kidd, C. E., Kidd, M. R. & Hofmann, H. A. Measuring multiple hormones from a single water sample using enzyme immunoassays. Gen. Comp. Endocrinol. 165, (2010).

32. Butler, J. M., Whitlow, S. M., Roberts, D. A. & Maruska, K. P. Neural and behavioural correlates of repeated social defeat. Sci. Rep. 8, 6818 (2018).

33. Knight, Z. A. et al. Resource Molecular Profiling of Activated Neurons by Phosphorylated Ribosome Capture. Cell 151, 1126–1137 (2012).

34. Maruska, K. P., Butler, J. M., Field, K. E., Forester, C. & Augustus, A. Neural Activation Patterns Associated with Maternal Mouthbrooding and Energetic State in an African Cichlid Fish. Neuroscience (2020) doi:10.1016/j.neuroscience.2020.07.025.

35. Fang, J. & Clemens, L. G. Contextual determinants of female–female mounting in laboratory rats. Anim. Behav. 57, 545–555 (1999).

36. Parker, G. A. Assessment strategy and the evolution of fighting behaviour. J. Theor. Biol. 47, 223–243 (1974).

37. Draud, M. Female and male Texas cichlids (Herichthys cyanoguttatum) do not fight by the same rules. Behav. Ecol. 15, 102–108 (2004).

38. Williams, D. I. & Wells, P. A. Differences in home-cage-emergence in the rat in relation to infantile handling. Psychon. Sci. 18, 168–169 (1970).

39. Oliveira, R. F., Carneiro, L. A. & Canário, A. V. M. No hormonal response in tied fights. Nature 437, 207–208 (2005).

40. Oliveira, R. F. Social behavior in context: Hormonal modulation of behavioral plasticity and social competence. Integr. Comp. Biol. 49, 423–440 (2009).

41. Oliveira, R. F. & Canário, A. V. M. Nemo through the looking-glass: a commentary on Desjardins & Fernald. Biol. Lett. 7, 487–488 (2011).

42. Hirschenhauser, K., Wittek, M., Johnston, P. & Möstl, E. Social context rather than behavioral output or winning modulates post-conflict testosterone responses in Japanese quail (Coturnix japonica). Physiol. Behav. 95, 457–463 (2008).

43. Loveland, J. L. & Fernald, R. D. Differential activation of vasotocin neurons in contexts that elicit aggression and courtship. Behav. Brain Res. 317, 188–203 (2017).

44. O’Connell, L. A., Rigney, M. M., Dykstra, D. W. & Hofmann, H. A. Neuroendocrine Mechanisms Underlying Sensory Integration of Social Signals. J. Neuroendocrinol. 25, 644–654 (2013).

45. Field, K. E., McVicker, C. T. & Maruska, K. P. Sexually-Relevant Visual and Chemosensory Signals Induce Distinct Behaviors and Neural Activation Patterns in the Social African Cichlid, Astatotilapia burtoni. Front. Behav. Neurosci. 12, 267 (2018).

46. Butler, J. M. & Maruska, K. P. The Mechanosensory Lateral Line System Mediates Activation of Socially-Relevant Brain Regions during Territorial Interactions. Front. Behav. Neurosci. 10, (2016).

47. Folgueira, M., Anadón, R. & Yáñez, J. Experimental study of the connections of the telencephalon in the rainbow trout ( Oncorhynchus mykiss ). II: Dorsal area and preoptic region: Telencephalic Connections in Trout. J. Comp. Neurol. 480, 204–233 (2004).

48. Yamamoto, N. & Ito, H. Fiber connections of the anterior preglomerular nucleus in cyprinids with notes on telencephalic connections of the preglomerular complex. J. Comp. Neurol. 491, 212–233 (2005).

49. Yamamoto, N. & Ito, H. Visual, lateral line, and auditory ascending pathways to the dorsal telencephalic area through the rostrolateral region of the lateral preglomerular nucleus in cyprinids. J. Comp. Neurol. 508, 615–647 (2008).

50. Desjardins, J. K., Klausner, J. Q. & Fernald, R. D. Female genomic response to mate information. Proc. Natl. Acad. Sci. 107, (2010).

51. Lin, D. et al. Functional identification of an aggression locus in the mouse hypothalamus. Nature 470, 221–226 (2011).

52. Yang, T. et al. Social Control of Hypothalamus-Mediated Male Aggression. Neuron 95, 955-970.e4 (2017).

53. Lin, D. et al. Functional identification of an aggression locus in the mouse hypothalamus. Nature 470, 221–226 (2011).

54. Maruska, K. P., Butler, J. M., Field, K. E. & Porter, D. T. Localization of glutamatergic, GABAergic, and cholinergic neurons in the brain of the African cichlid fish, Astatotilapia burtoni. J. Comp. Neurol. 525, 610–638 (2017).

55. Lee, H. et al. Scalable control of mounting and attack by Esr1+ neurons in the ventromedial hypothalamus. Nature 509, 627–632 (2014).

