## Supplementary Material for "Sex differences in aggression and its neural substrate in a cichlid fish"

### Supplementary Results

#### *Statistical results of comparisons between NC and mirror fish for behaviors*

There was a significant effect of condition but not sex or an interaction on lateral displays type 1 (Table S1; lateral displays type 1, Two-way ANOVA, condition: p = 0.1856). There was a significant effect of sex, condition, and sex*condition on lateral displays type 2 (Table S1; lateral displays type 2, Two-way ANOVA, sex: p = 0.0480, condition: p = 0.0137, interaction: p = 0.0480). Post-hoc Šídák's tests reveal that males in the mirror condition perform more type 1 and type 2 lateral displays than negative control males, but females do not differ by condition (Table S1; lateral displays type 1: p = 0.0112, lateral displays type 2: p = 0.0092, Šídák's tests). Furthermore, there was a significant effect of condition, but not sex or an interaction on s-curve displays, mouth contact, and taps (Table S1; s-curve displays, Two-way ANOVA, condition: p = 0.0072, Table S1; mouth contact, Two-way ANOVA of log transformation, condition: p < 0.0001, Table S1; taps, Two-way ANOVA of log transformation, condition: p = 0.0031). Mirror males perform more s-curve displays than negative control males, but females do not differ by condition (Table S1; s-curve displays, Šídák's test, Male Mirror – NC, p = 0.0078). Both mirror males and females perform more mouth contact behaviors than negative controls (Table S1; mouth contact, Male Mirror – NC: p < 0.0001, Female Mirror – NC: p < 0.0001, Šídák's test). Mirror females perform more taps than negative control females while males do not differ by condition (Table S1; taps, Šídák's test, p = 0.0198).

There was a significant effect of condition but not sex or an interaction on total number of behaviors and total number of aggressive behaviors (Table S1; # of behaviors, Two-way ANOVA of log transformation, condition: p < 0.0001, Table S1; total aggression, Two-way ANOVA, condition: p < 0.0001). Both mirror males and mirror females perform more total behaviors and aggressive behaviors than negative controls (Table S1; # of behaviors, Male Mirror – NC: p < 0.0001, Female Mirror – NC: p < 0.0001; total aggression, Male Mirror – NC: p = 0.0050, Female Mirror – NC: p = 0.0033, Šídák's tests).

1

| | Sex | | Condition | | Sex*Condition | |
|---|---|---|---|---|---|---|
| | F | P | F | P | F | P |
| Lateral Display Type 1 | 1.849<br>1, 26 | 0.1856 | 10.49<br>1, 26 | **0.0033** | 1.849<br>1, 26 | 0.1856 |
| Lateral Display Type 2 | 4.307<br>1, 26 | **0.0480** | 6.995<br>1, 26 | **0.0137** | 4.307<br>1, 26 | **0.0480** |
| S-curve Displays | 3.637<br>1, 26 | 0.0676 | 8.501<br>1, 26 | **0.0072** | 3.637<br>1, 26 | 0.0676 |

| | | | | | | |
|---|---|---|---|---|---|---|
| Mouth Contact | 0.6808<br>1, 26 | 0.4168 | 67.36<br>1, 26 | **< 0.0001** | 0.2122<br>1, 26 | 0.6489 |
| Taps | 0.1947<br>1, 26 | 0.6627 | 10.63<br>1, 26 | **0.0031** | 0.1218<br>1, 26 | 0.7299 |
| # of Behaviors | 0.4261<br>1, 26 | 0.5197 | 88.75<br>1, 26 | **< 0.0001** | 0.01433<br>1, 26 | 0.9056 |
| Total Aggression | 0.03252<br>1, 26 | 0.8583 | 23.23<br>1, 26 | **< 0.0001** | 0.07211<br>1, 26 | 0.7904 |

2

| | Male Mirror – NC | Female Mirror – NC |
|---|---|---|
| | P | P |
| Lateral Display Type 1 | **0.0112** | 0.2935 |
| Lateral Display Type 2 | **0.0092** | 0.8873 |
| S-curve Displays | **0.0078** | 0.6908 |
| Mouth Contact | **< 0.0001** | **< 0.0001** |
| Taps | 0.1293 | **0.0198** |
| # of Behaviors | < 0.0001 | < 0.0001 |
| Total Aggression | 0.0050 | 0.0033 |

Table S1. (1) Effect of sex and condition on behaviors quantified in a mirror assay. (2) Post-hoc comparisons.

| | Sex | | Condition | | Sex*Condition | |
|---|---|---|---|---|---|---|
| | F | P | F | P | F | P |
| POA | 0.8266<br>1, 25 | 0.3719 | 2.494<br>1, 25 | 0.1268 | 0.003165<br>1, 25 | 0.9556 |
| nPPa | 2.210<br>1, 25 | 0.1497 | 2.744<br>1, 25 | 0.1101 | 0.01917<br>1, 25 | 0.8910 |
| nPMp | 0.6952<br>1,19 | 0.4148 | 0.4276<br>1, 19 | 0.5210 | 0.3621<br>1, 19 | 0.5545 |
| Dl-g | 2.497<br>1, 24 | 0.1271 | 0.03271<br>1, 24 | 0.8580 | 1.146<br>1, 24 | 0.2951 |
| Dm-3 | 0.7580<br>1, 25 | 0.3922 | 0.4440<br>1, 25 | 0.5113 | 3.073<br>1, 25 | 0.0919 |
| Xm | 2.314<br>1, 23 | 0.1418 | 2.800<br>1, 23 | 0.1078 | 0.003933<br>1, 23 | 0.9505 |

Table S2. Effect of sex and condition on pS6 immunoreactivity.

| | | Vs | ATn | PAG | CG | nPPa | nMMp | nPMp | POA | VL | Xm | Dm-3 | Dl-g |
|---|---|---|---|---|---|---|---|---|---|---|---|---|---|
| Vv | R | -0.07 | -0.29 | -0.01 | 0 | 0.42 | 0.27 | 0.35 | 0.36 | -0.01 | **0.6** | -0.41 | 0.34 |
| | P | 0.8251 | 0.3391 | 0.9777 | 0.9966 | 0.1481 | 0.3802 | 0.2444 | 0.225 | 0.9869 | **0.0291** | 0.1662 | 0.2579 |
| Vs | R | | -0.14 | -0.48 | **0.75** | **0.58** | 0.5 | **0.61** | 0.54 | -0.28 | -0.34 | **-0.59** | 0.24 |
| | P | | 0.6569 | 0.0943 | **0.0031** | **0.037** | 0.0788 | **0.0262** | 0.0553 | 0.3466 | 0.2606 | **0.0336** | 0.4221 |
| ATn | R | | | -0.33 | -0.43 | 0.11 | 0.48 | 0.16 | 0.3 | -0.2 | 0.21 | -0.25 | -0.48 |
| | P | | | 0.2674 | 0.146 | 0.7209 | 0.0957 | 0.5941 | 0.3156 | 0.5184 | 0.4968 | 0.4067 | 0.0939 |
| PAG | R | | | | 0.16 | -0.25 | -0.44 | -0.39 | -0.37 | 0.08 | -0.15 | **0.57** | -0.15 |
| | P | | | | 0.609 | 0.4008 | 0.1338 | 0.1825 | 0.2085 | 0.8064 | 0.6306 | **0.041** | 0.6188 |
| CG | R | | | | | 0.47 | 0.23 | 0.4 | 0.34 | -0.22 | -0.47 | -0.26 | 0.27 |
| | P | | | | | 0.1034 | 0.45 | 0.1721 | 0.2594 | 0.4777 | 0.1045 | 0.392 | 0.375 |
| nPPa | R | | | | | | **0.89** | **0.94** | **0.97** | **-0.73** | 0.24 | **-0.84** | -0.13 |
| | P | | | | | | **<0.001** | **<0.001** | **<0.001** | **0.0048** | 0.4221 | **0.0003** | 0.6669 |
| nMMp | R | | | | | | | **0.93** | **0.97** | **-0.73** | 0.4 | **-0.92** | -0.33 |
| | P | | | | | | | **<0.001** | **<0.001** | **0.0049** | 0.1808 | **<0.001** | 0.2697 |
| nPMp | R | | | | | | | | **0.97** | **-0.79** | 0.37 | **-0.92** | -0.25 |
| | P | | | | | | | | **<0.001** | **0.0013** | 0.2116 | **<0.001** | 0.4082 |
| POA | R | | | | | | | | | **-0.76** | 0.35 | **-0.91** | -0.24 |
| | P | | | | | | | | | **0.0023** | 0.2382 | **<0.001** | 0.4235 |
| VL | R | | | | | | | | | | -0.31 | **0.63** | **0.62** |
| | P | | | | | | | | | | 0.305 | **0.0218** | **0.0248** |
| Xm | R | | | | | | | | | | | -0.49 | -0.4 |
| | P | | | | | | | | | | | 0.0924 | 0.1803 |
| Dm-3 | R | | | | | | | | | | | | 0.13 |
| | P | | | | | | | | | | | | 0.6712 |

Table S3. Pearson correlation values of pS6 immunoreactivity in mirror fish.

| | | Vs | ATn | PAG | CG | nPPa | nMMp | nPMp | POA | VL | Xm | Dm-3 | Dl-g |
|---|---|---|---|---|---|---|---|---|---|---|---|---|---|
| Vv | R | **0.79** | **0.96** | -0.18 | **-0.59** | 0.11 | -0.06 | **0.59** | 0.01 | -0.39 | -0.42 | **0.76** | 0.42 |
| | P | **0.0013** | **<0.001** | 0.5522 | **0.0357** | 0.7121 | 0.8519 | **0.0344** | 0.9646 | 0.1871 | 0.1517 | **0.0024** | 0.1481 |
| Vs | R | | **0.73** | **-0.62** | -0.2 | 0.36 | -0.19 | **0.6** | 0.13 | 0.21 | 0.19 | **0.95** | 0.39 |
| | P | | **0.0043** | **0.0245** | 0.5213 | 0.2292 | 0.5235 | **0.0307** | 0.6796 | 0.5 | 0.5351 | **<0.001** | 0.1869 |
| ATn | R | | | -0.12 | -0.52 | 0.18 | -0.05 | **0.56** | 0.04 | -0.46 | -0.44 | **0.67** | 0.34 |
| | P | | | 0.7081 | 0.066 | 0.5542 | 0.8606 | **0.0486** | 0.9034 | 0.112 | 0.1361 | **0.0117** | 0.262 |
| PAG | R | | | | -0.38 | -0.12 | 0.46 | -0.11 | 0.1 | **-0.73** | **-0.61** | **-0.59** | 0.17 |
| | P | | | | 0.1993 | 0.6908 | 0.1148 | 0.7208 | 0.7522 | **0.0049** | **0.026** | **0.0336** | 0.5858 |
| CG | R | | | | | -0.15 | -0.42 | -0.45 | -0.26 | **0.66** | **0.63** | -0.32 | -0.46 |
| | P | | | | | 0.6293 | 0.1561 | 0.126 | 0.3958 | **0.0137** | **0.0217** | 0.2821 | 0.1157 |
| nPPa | R | | | | | | 0.18 | 0.43 | **0.64** | 0.25 | 0.42 | 0.48 | 0.16 |
| | P | | | | | | 0.5622 | 0.1453 | **0.0179** | 0.4176 | 0.1578 | 0.0951 | 0.6016 |
| nMMp | R | | | | | | | **0.62** | **0.83** | -0.28 | -0.34 | -0.15 | -0.17 |
| | P | | | | | | | **0.0224** | **0.0005** | 0.3608 | 0.2591 | 0.63 | 0.5867 |
| nPMp | R | | | | | | | | **0.77** | -0.09 | -0.2 | **0.62** | 0.02 |
| | P | | | | | | | | **0.0022** | 0.7593 | 0.523 | **0.0239** | 0.9374 |
| POA | R | | | | | | | | | 0.08 | 0.06 | 0.21 | -0.17 |
| | P | | | | | | | | | 0.7908 | 0.8363 | 0.4857 | 0.5736 |
| VL | R | | | | | | | | | | **0.93** | 0.23 | -0.15 |
| | P | | | | | | | | | | **<0.001** | 0.4573 | 0.6326 |
| Xm | R | | | | | | | | | | | 0.2 | 0.01 |
| | P | | | | | | | | | | | 0.5127 | 0.9715 |
| Dm-3 | R | | | | | | | | | | | | 0.41 |
| | P | | | | | | | | | | | | 0.1596 |

Table S4. Pearson correlation values of pS6 immunoreactivity in negative control fish.

| | PC1 | PC2 |
|---|---|---|
| Vv | 0.22776301 | 0.26646841 |
| Vs | 0.23783291 | 0.24196108 |
| ATn | 0.10442256 | -0.2176296 |
| PAG | 0.00044817 | 0.24869525 |
| CG | 0.20889062 | **0.42034068** |
| nPPa | **0.40851882** | 0.06593241 |
| nMMp | **0.42541373** | -0.1225907 |
| nPMp | **0.41868272** | -0.046673 |
| POA | **0.44725236** | -0.0550992 |
| VL | -0.0798084 | **0.43327604** |
| Xm | 0.12684742 | -0.2566584 |
| Dm-3 | -0.2984527 | 0.2373274 |
| Dl-g | 0.04787439 | **0.50003537** |

Table S5. PCA component loadings for mirror fish.

| | PC1 | PC2 |
|---|---|---|
| Vv | 0.2776275 | **-0.3914845** |
| Vs | 0.322159 | -0.068534 |
| ATn | 0.2796261 | **-0.3441852** |
| PAG | 0.1820699 | -0.1765652 |
| CG | 0.2067307 | **0.42871937** |
| nPPa | 0.2962492 | 0.07486987 |
| nMMp | 0.2415449 | -0.1042493 |
| nPMp | **0.3299453** | -0.1190802 |
| POA | **0.3188154** | 0.02304585 |
| VL | 0.251112 | **0.49804965** |
| Xm | 0.2684649 | **0.46637798** |
| Dm-3 | **0.306559** | -0.090484 |
| Dl-g | 0.2816647 | -0.0605815 |

Figure S6. PCA component loadings for negative control fish.

**Supplementary Figures**

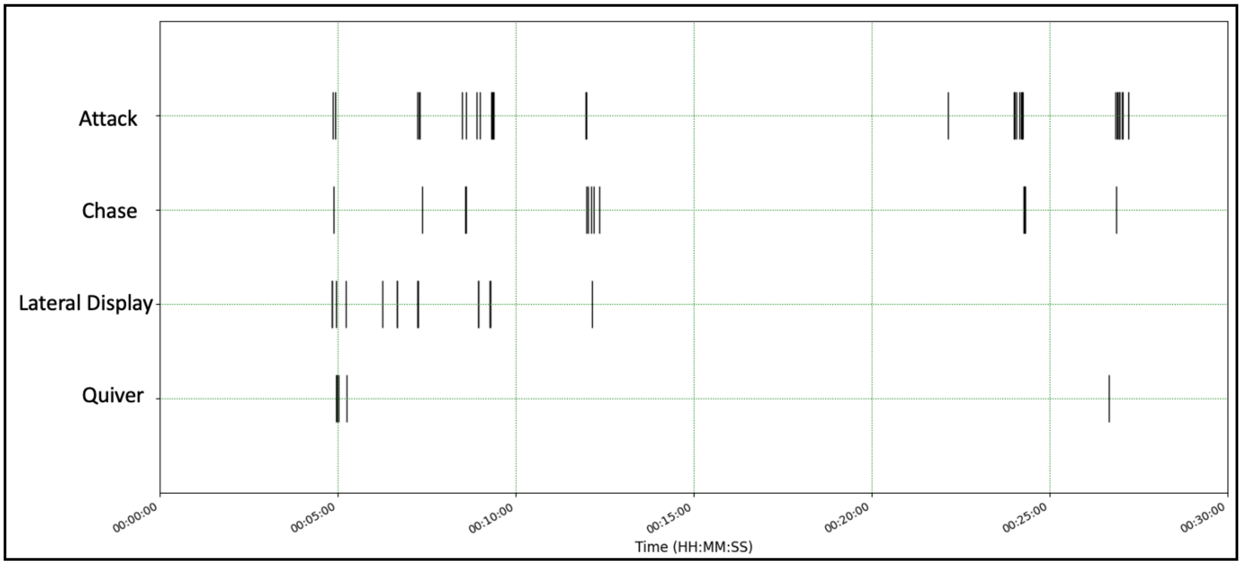


**Figure S1. Representative raster plot of aggressive behaviors performed by a focal female in a dyad assay.** Gravid females perform aggressive behaviors (including quivers) in an assay with a live opponent. The first 30 minutes of the behaviors performed in a dyad assay are represented here.

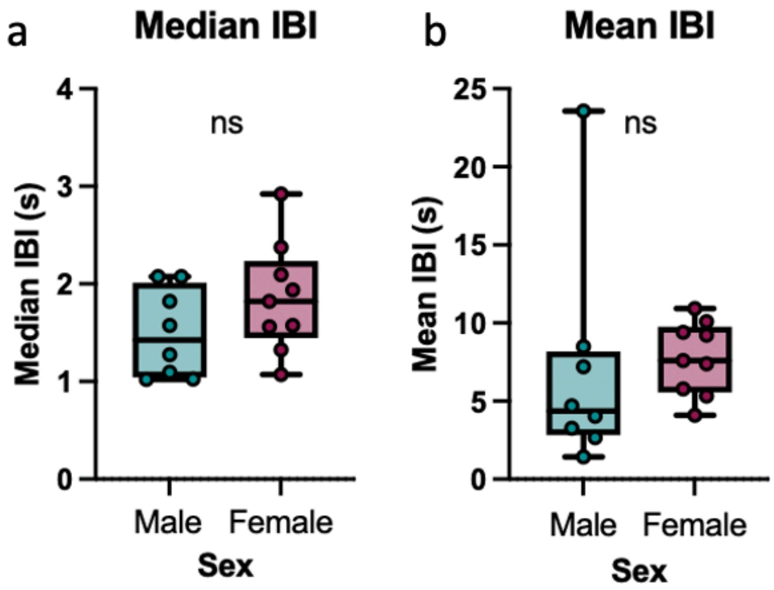


**Figure S2. No sex differences in inter-behavioral intervals.** (a)(b) Mirror males and females do not differ by median and mean inter-behavioral intervals.

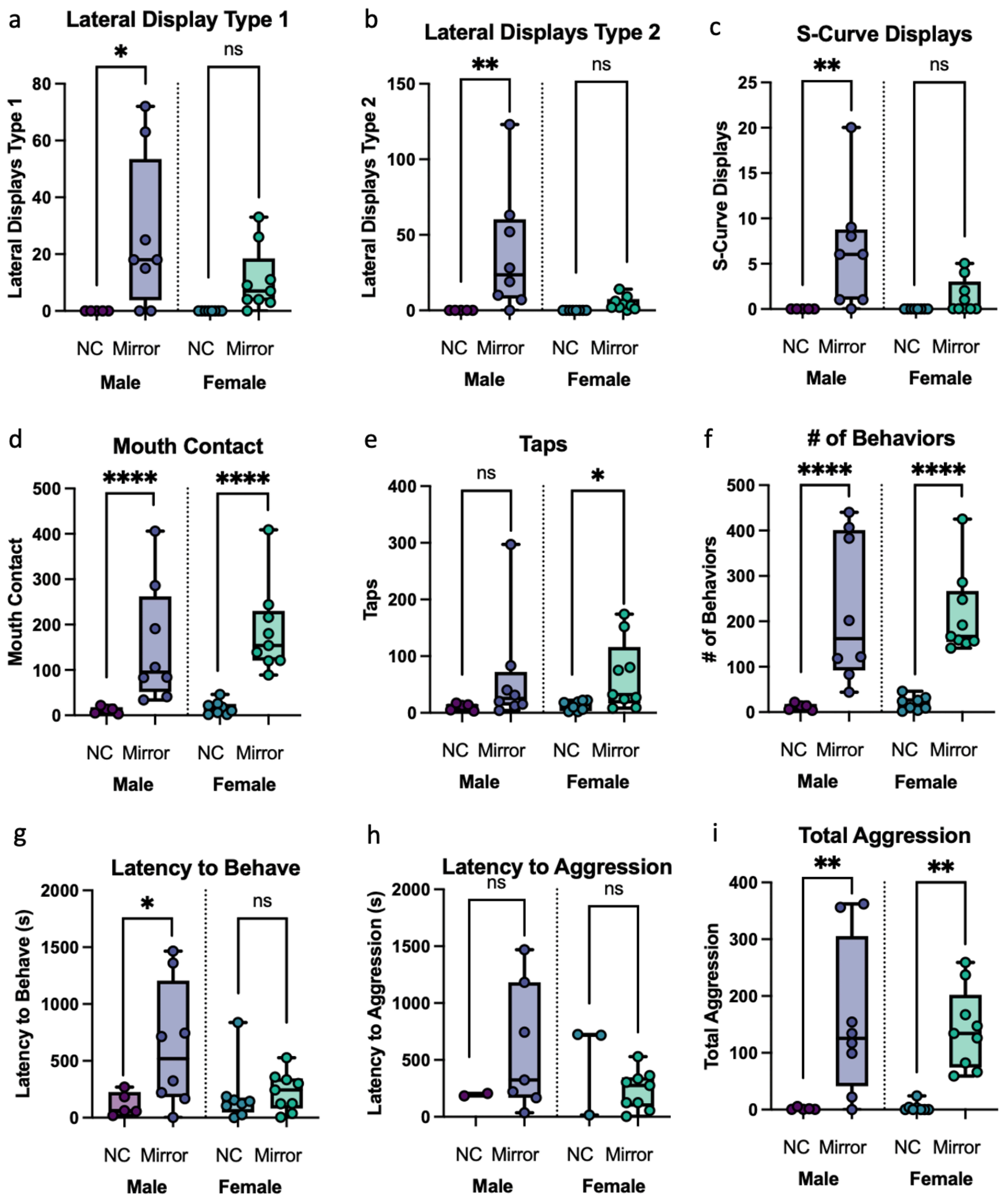


**Figure S3. *A. burtoni* perform more behaviors when exposed to a mirror compared to an opaque cover.** (a)(b) Mirror males perform more lateral display type 1 and 2 than negative control males while females do not differ by condition. (c) Mirror males perform more s-curve displays than negative control males while females do not differ by condition. (d) Mirror males and females perform more mouth contact behaviors than negative control fish. (e) Mirror females perform more taps than negative control females while males do not differ by condition. (f) Mirror males and females perform more behaviors than negative control fish. (g) Mirror males have a longer latency to behave than negative control males while females do not differ by condition. (h) The latency to aggression does not statistically differ by sex or condition. (i) Mirror males and females perform more aggressive behaviors than negative control fish.

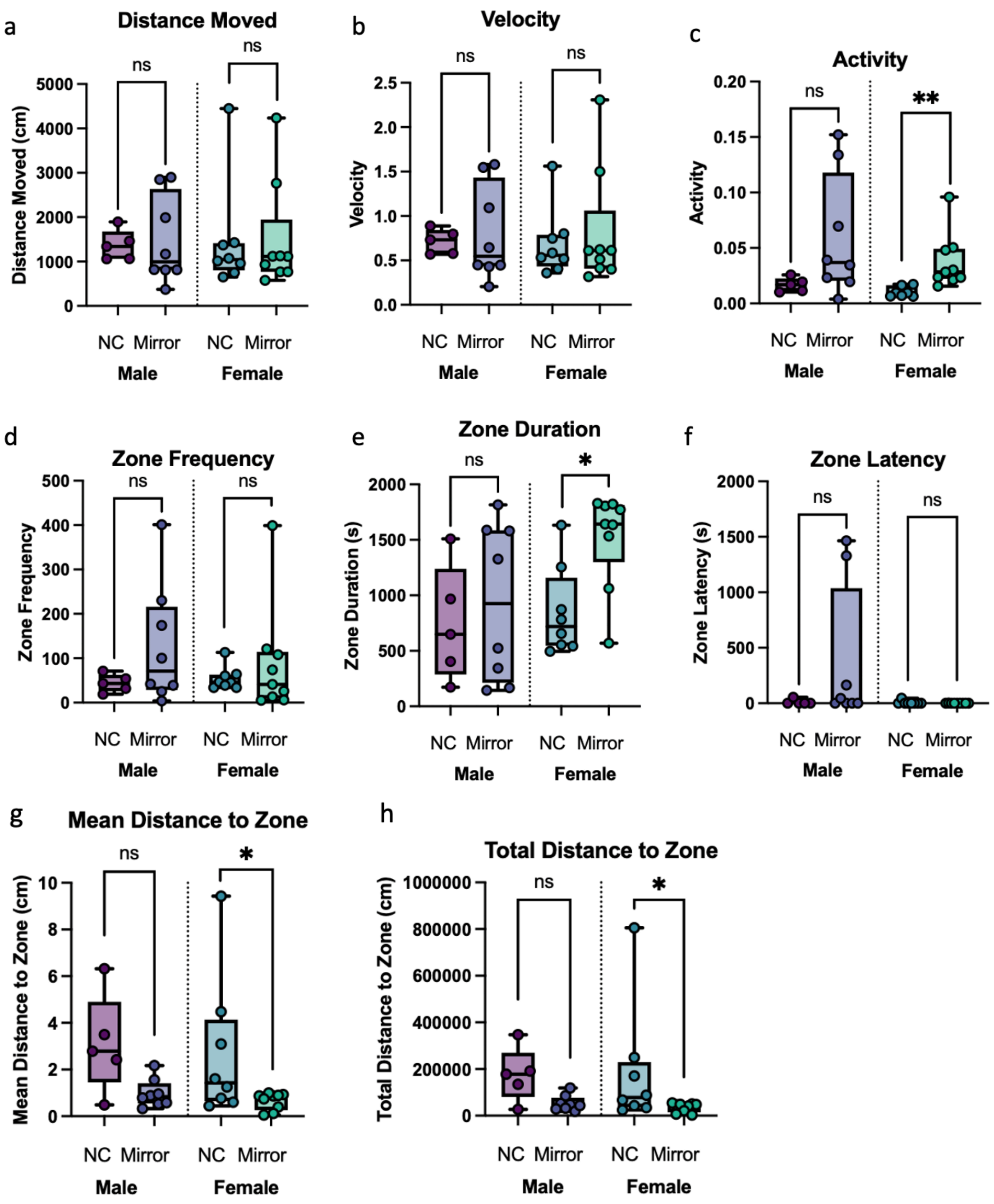


**Figure S4. Similarities and differences in Noldus tracking data in fish exposed to a mirror or opaque cover.** (a)(b) Distance moved and velocity did not differ by sex or condition. (c) Mirror females were more active than negative control females. Mirror males followed this pattern although it was not statistically significant. (d) Zone frequency did not differ by sex or condition. (e) Mirror females spent more time in the zone than negative control females while males did not differ by condition. (f) Zone latency did not differ by sex or condition. (g)(h) Negative control females maintained a larger mean and total distance from the zone than negative control females. Negative control males followed this pattern although it was not statistically significant.

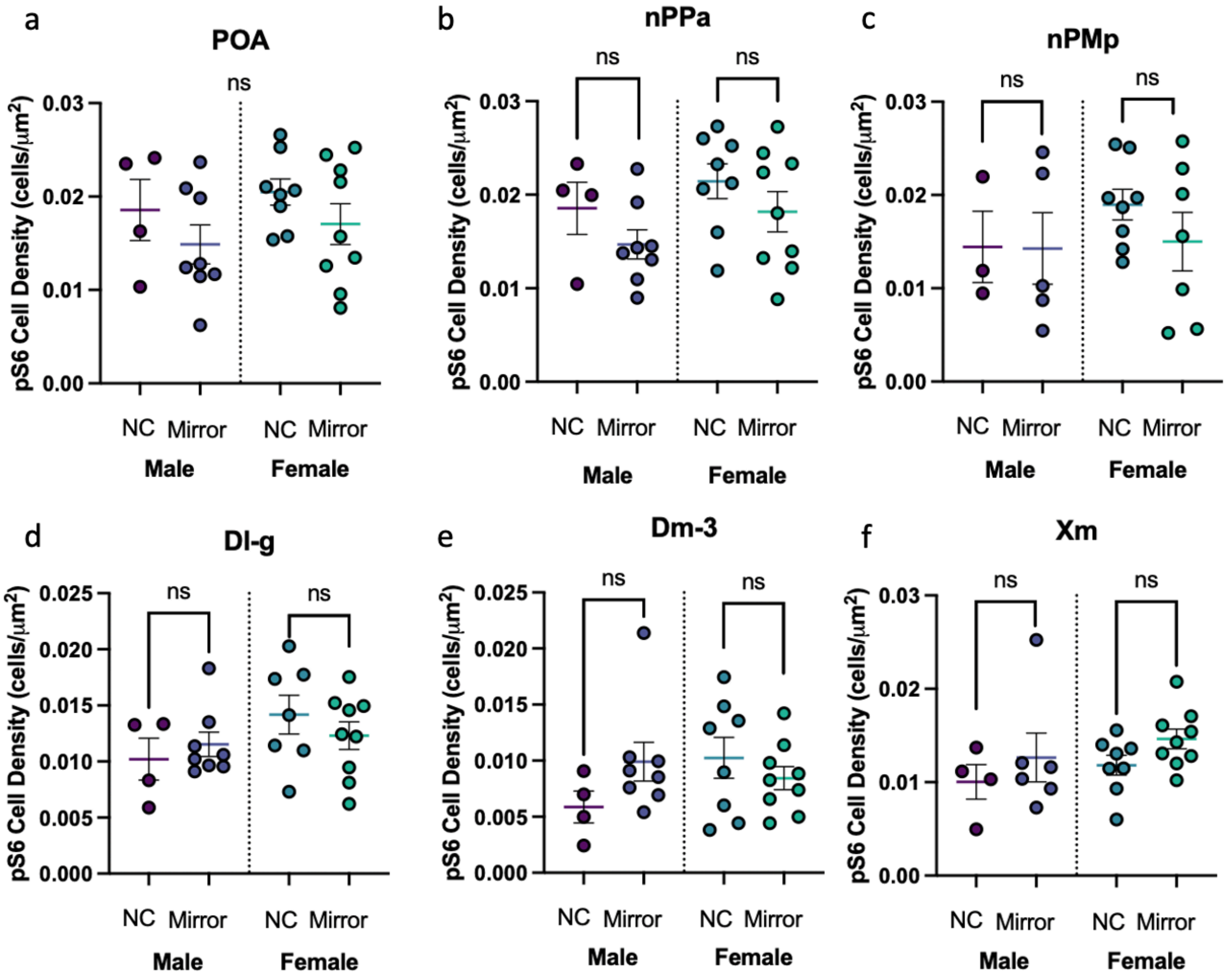


**Figure S5. pS6 immunoreactivity in brain regions relevant to social behaviors and sensory processing.** (a) pS6 cell density does not differ by condition or sex in the POA. (b)(c)(d)(e)(f) pS6 immunoreactivity did not differ by sex or condition in the nPPa, nPMp, Dl-g, Dm-3, or Xm.

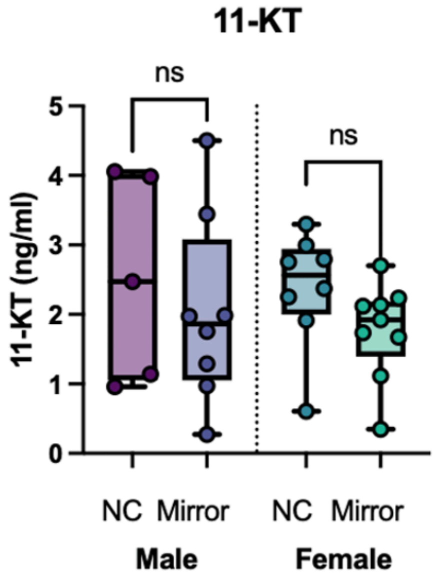


**Figure S6. Physiological measures between sex and condition.** 11-KT did not differ between sex and condition.

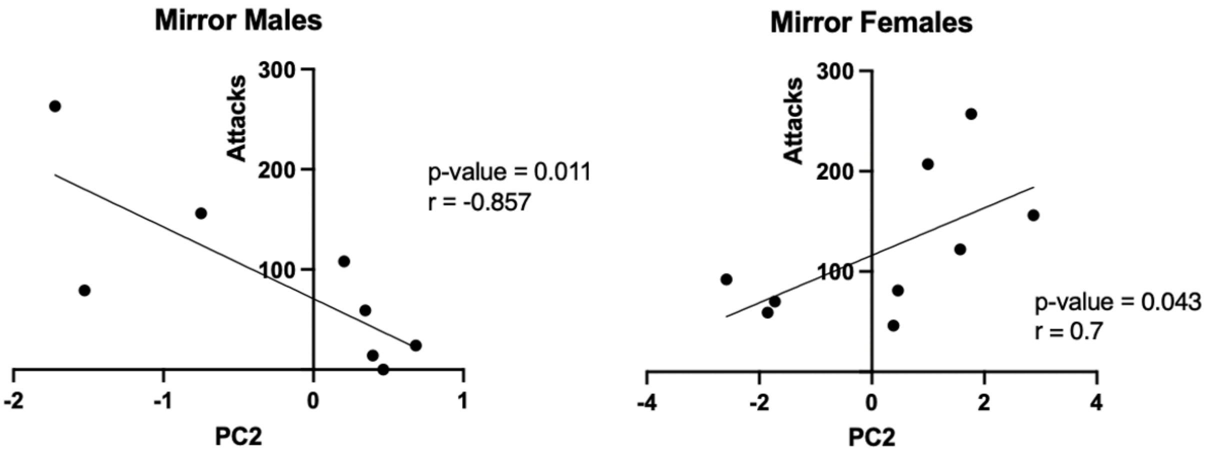


**Figure S7. Correlations of PC2 to attacks for mirror males and mirror females.**
